# An ancient genome duplication in the speciose reef-building coral genus, *Acropora*

**DOI:** 10.1101/366435

**Authors:** Yafei Mao, Chuya Shinzato, Noriyuki Satoh

**Author notes:** Present address: Atmosphere and Ocean Research Institute, The University of Tokyo, Kashiwa, Chiba 277-8564, Japan (Chuya Shinzato). Correspondence to: Yafei Mao.

## Abstract

Whole-genome duplication (WGD) has been recognized as a significant evolutionary force in the origin and diversification of vertebrates, plants, and other organisms. *Acropora*, one of the most speciose reef-building coral genera, responsible for creating spectacular but increasingly threatened marine ecosystems, is suspected to have originated by polyploidy, yet there is no genetic evidence to support this hypothesis. Using comprehensive phylogenomic and comparative genomic approaches, we analyzed five *Acropora* genomes and an *Astreopora* genome (Scleractinia: Acroporidae) to show that a WGD event likely occurred between 27.9 and 35.7 Million years ago (Mya) in the most recent common ancestor of *Acropora*, concurrent with a massive worldwide coral extinction. We found that duplicated genes became highly enriched in gene regulation functions, some of which are involved in stress responses. The different functional clusters of duplicated genes are related to the divergence of gene expression patterns during development. Some gene duplications of proteinaceous toxins were generated by WGD in *Acropora* compared with other Cnidarian species. Collectively, this study provides evidence for an ancient WGD event in corals and it helps to explain the origin and diversification of *Acropora*.

## Introduction

Reef-building corals contribute to tropical marine ecosystems that support innumerable marine organisms, but reefs are increasingly threatened due to recent increases in seawater temperatures, pollution, and other stressors [1,2]. The Acroporidae is a family of reef-building corals in the phylum Cnidaria, one of the basal phyla of the animal clade [3-5]. *Astreopora* (Anthozoa: Acroporidae) is the sister genus in the acroporid lineage according to fossil records and molecular phylogenetic evidence [3,6,7]. Importantly, *Acropora* (Anthozoa: Acroporidae), one of the most diverse genera of reef-building corals, including more than 150 species in Indo-Pacific Ocean, is thought to have originated from *Astreopora* almost 60 Mya with several species turnovers [3,5,8,9]. Investigating the evolutionary history of this group importantly contributes to our understanding of coral reef biodiversity and conservation. Hybridization among *Acropora* species has been observed in the wild [10] and variable chromosome numbers have been determined in different *Acropora* lineages [11]. Additionally, gene duplications have been shown in several *Acropora* gene families [12,13]. Thus, based on their unique lifestyle, variable chromosome numbers, and complicated reticular evolutionary history, Indo-Pacific *Acropora* likely originated via polyploidy [10-16]. However, there is no direct molecular and genetic evidence to support this hypothesis.

Ancient whole (large-scale)-genome duplication (WGD), or paleopolyploidy, has shaped in the genomes of vertebrates, green plants, and other organisms, and is usually regarded as an evolutionary landmark in the origins and diversification of organisms [17-19] (Supplementary Fig. 1). Two separate WGD events have been documented in the common ancestors of vertebrates (two-rounds of WGD) [20] and another major WGD has been reported in the last common ancestor of teleost fish [21,22]. Meanwhile, living angiosperms share an ancient WGD event [23,24], and many other WGD events have been reported in major clades of angiosperms [25,26]. In addition, two-rounds of WGDs in the vertebrates are suggested to have occurred during the Cambrian Period, and some WGDs in plants are believed to have occurred during Cretaceous-Tertiary [17,25,27]. Thus, WGD is regarded as an important evolutionary way to reduce the risk of extinction [17,18,25]. However, the study of WGD in Cnidaria has received less attention [17,18,28-30].

**Figure 1.**
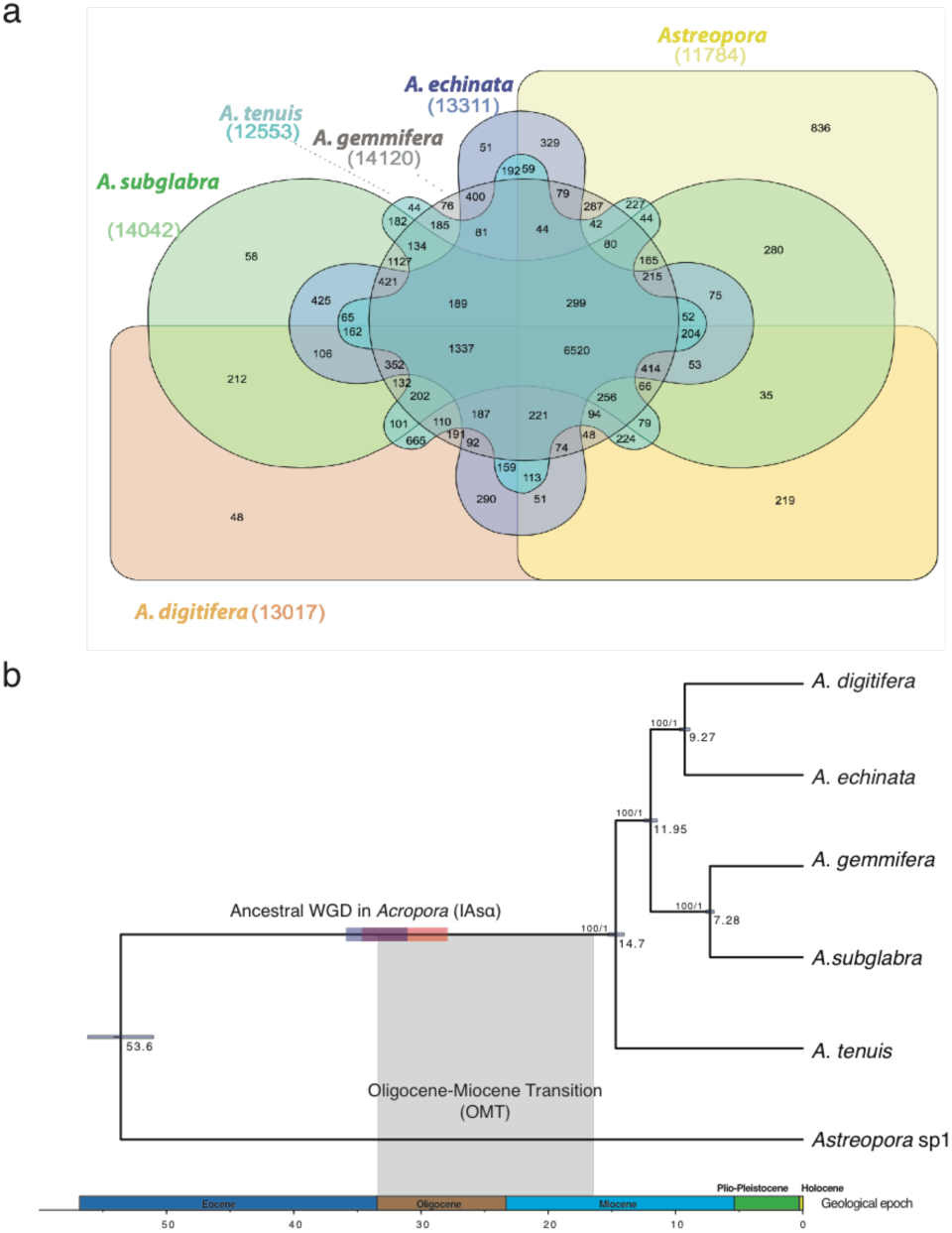
Ancient WGD in the reef-building coral *Acropora* (IAsα). (a). Venn diagram of shared and unique gene families in six *Acroporid* species. (b). A calibrated phylogenomic tree of six *Acroporid* species inferred from 3,461 single-copy orthologs using BEAST2. Horizontal bars on branches of the tree represent the timing of WGD in *Acropora*. The timing of IAsα was estimated at 35.01 Mya (95% confidence interval: 31.18-35.7 Mya) by dS-based analysis (horizontal blue bar) and 30.78 Mya (95% confidence interval: 27.86-34.77 Mya) by phylogenomic analysis (horizontal orange bar). Grey shading represents the timing of one coral species turnover event, the Oligocene-Miocene transition (OMT), suggesting that IAsα is correlated with OMT.

Duplicated genes created by WGD have complex fates following the time of diploidization [18,31]. Usually, one of the duplicated genes is silenced or lost due to redundancy of gene functions, termed “nonfunctionalization”. However, retained duplicated genes provide important sources of biological complexity and evolutionary novelty due to subfunctionalization, neofunctionalization, and dosage effects [22,32]. Duplicated genes may develop complementary gene functions via subfunctionalization, or evolve new functions through neofunctionalization, or are retained in complicated regulatory networks with different gene expressions due to dosage effects. For instance, duplicated MADS-Box genes are crucial for flower development and the origin of phenotypic novelty in plants [18,33]. Duplicated homeobox genes provide raw genetic material for vertebrate development [22,34]. In addition, toxin diversification following by gene duplications has been recognized as a mechanism to enhance adaptation in animals [35,36], especially in snake venoms [37,38]. Meanwhile, toxic proteins are involved in various important processes in corals, including prey capture, protection from predators, wound-healing, etc. [39,40], but it is still unclear how gene duplications of toxic proteins evolved in corals.

Isozyme electrophoresis and restriction fragment length polymorphism (RFLP) were used to identify gene duplications in polyploids a few decades ago [41,42]. In the past ten years, next generation sequencing (NGS) has generated a wealth of genomic data at vastly decreased cost and reduced effort [43,44]. Then, three main methods were developed to identify WGD, based on: 1). analysis of the rate of synonymous substitutions per synonymous site (dS) of duplicated genes within a genome (dS-based method) [25,45,46]; 2). phylogenetic analysis of gene families among multiple genomes (Phylogenomic analysis) [23,47]; and 3). synteny block identification compared with sister lineages without WGD (Synteny analysis) [20,48,49], respectively. The dS-based method and phylogenomic analysis only require gene family information, without genome assembly. However, the dS-based method cannot detect ancient WGD, and gene tree uncertainty usually causes bias in the phylogenomic analysis. Both methods rely heavily on gene family estimation and clustering. Inaccurate gene predictions (gene models) and rough gene family cluster algorithms can easily fail to detect WGD using either method. In contrast, the synteny analysis relies heavily on the genome assembly quality. Poor assembly quality can hide the WGD signals, and some genomes with huge rearrangements cannot be used to detect WGD using synteny block identification. Thus, the most credible conclusions depend on complementary evidence from different methods [24,50,51].

Here, we analyzed a genome of *Astreopora* (*Astreopora* sp1) as an outgroup, as well as five *Acropora* genomes (*A. digitifera, A. gemmifera, A. subgrabla, A. echinata* and *A. tenuis*) to address the following questions using all three methods; (I) whether and when WGD occurred in *Acropora*, (II) what is the fate of duplicated genes in *Acropora* after the event, (III) what the gene expression patterns of duplicated genes cross five developmental stages in *A. ditigifera*, and (IV) what roles of WGD were involved in the diversification of toxic proteins in *Acropora* (Supplementary Fig. 2).

## Results

### Cluster of gene families and calibration of the acroporid phylogenomic tree

We clustered all homologs among the six *Acroporid* species into 19,760 gene families, and they shared 6,520 gene families (Fig 1a, See Additional file 1). Each *Acropora* genome had very few unique gene families, but *Astreopora* sp1 had 218, suggesting that *Astreopora* sp1 is genetically divergent from the five *Acropora* species. 3,461 single-copy orthologs were selected from 6,520 shared gene families. These were concatenated to reconstruct a calibrated phylogenomic tree based on the reported divergence time of *Acropora* (Mao et al., in revision). We found that *Astreopora* sp1 split from *Acropora* ~ 53.6 Mya (95% highest posterior density (HPD): 51.02 - 56.21 My) (Fig 1b; Supplementary Fig 3). This established a timescale to analyze the timing of the subsequent WGD.

### WGD identification with the dS-based method

Synonymous substitution rate (dS) analysis has been widely used to infer WGD [25,52]. We identified over 10,000 paralogous gene pairs, based on their sequence similarities as well as we identified over 10,000 anchor gene pairs, based on synteny information from each species (Supplementary Table 1; for details see Methods). Then we calculated dS values from paralogous gene pairs and anchor gene pairs for each species.

An ‘L-shaped’ distribution was evident in both paralogous and anchor gene pair dS distributions of *Astreaopora* sp1, illustrating that no WGD occurred in *Astreaopora* sp1. However, all five *Acropora* species displayed a similar peak in dS distributions of both paralogous and anchor gene pairs (peak: 0~0.3), suggesting that WGD did occur in *Acropora* (Fig 2a, Supplementary Figs 4, 5).

**Figure 2.**
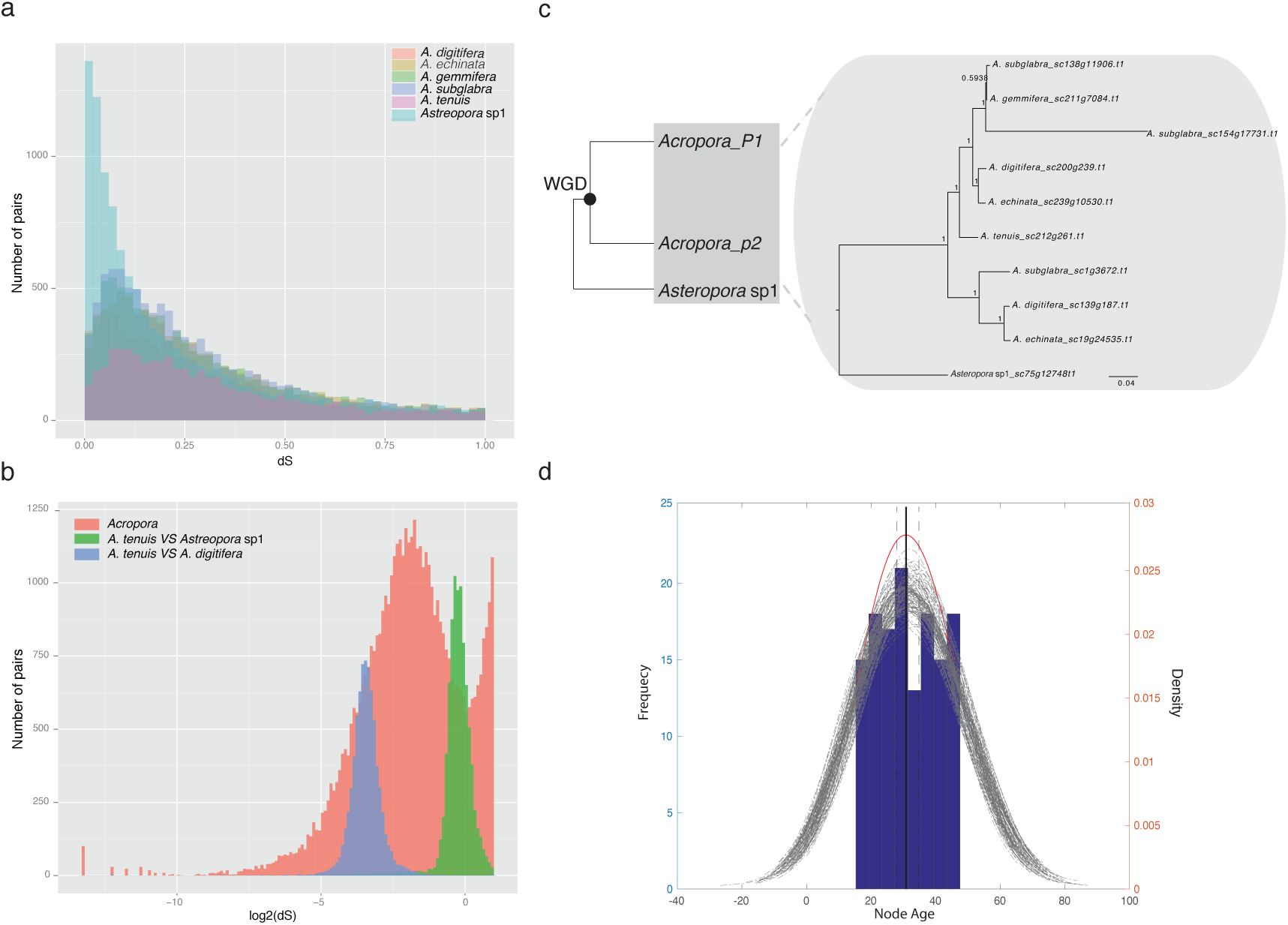
Ancient WGD identification (IAsα) and timing of the event in *Acropora*. (a). Frequency distribution of dS values for paralogous gene pairs in five *Acropora* and one *Astreopora* species showing that a WGD event occurred in *Acropora*. Similar peaks (dS value: 0.1-0.3) in dS distributions of five *Acropora* lineages indicate that a WGD event occurred in *Acropora*. (Light red: *A. digitifera*; Light yellow: *A. echinata*; Light green: *A. gmmifera*; Light blue: *A. subglabra*; Light purple: *A. tenuis*; Light cyan: *Astreopora* sp1). (b). Frequency distribution of dS log values for paralogous genes in *Acropora* and for orthologous genes showing that a WGD event occurred in the most recent common ancestor of *Acropora*. Distributions are plotted with a bin size of 0.1. (c). Hypothetical tree topology of duplicated genes in the *Acroporidae* and the phylogeny of one duplicated gene (alpha-protein kinase 1-like). The phylogenetic tree shows gene retention, loss, and duplications following with WGD. (d). Node age distribution of IAsα. Inferred node ages from 135 phylogenies were analyzed with KDE toolbox to show the peak at 30.78 My, represented by the black solid line. The grey lines represent density estimations from 1000 bootstraps and the black dotted line represents the corresponding 95% confidence interval (27.86 - 34.77 My) from 100 bootstraps.

dS values of orthologous gene pairs between two pairs of species (*Astreopora* sp1 and *A. tenuis; A. tenuis* and *A. digitifera*) were estimated as the speciation time between them according to neutral evolution theory [48,53]. We combined the dS values of paralogous gene pairs for the five *Acropora* species and estimated the peak in the log dS distribution (modal value = −1.82). Also, we estimated the distribution of orthologous gene pairs between *Astreopora* sp1 and *A. tenuis* (modal value = −0.31) and the distribution of orthologous gene pairs between *A. tenuis* and *A. digitifera* (modal value = −3.46). The result indicates that the WGD occurred in *Acropora* after the split of *Astreopora* sp1 and *A. tenuis* (Fig 2b). In other words, an ancient WGD event likely occurred in the most recent common ancestor of *Acropora*. Based on speciation time estimated in the calibrated phylogenomic tree and assuming a constant dS rate [25], we estimated that the WGD of *Acropora* occurred ~ 35.01 Mya (95% confidence interval: 31.18 - 35.7 My) (Supplementary Table 2, for details see Methods). Here, we defined this event as invertebrate α event of WGD specifically in *Acropora* (IAsα).

### Phylogenomic and synteny analysis of IAsα

If the proposed IAsα is correct, then the ohnologs of *Acropora* (paralogs created by IAsα) should form two clades from their orthologs in *Astreopora* sp1 by mapping IAsα onto phylogenetic trees [23,54]. In other words, the phylogenetic topology would be (((*Acropora clade1*) bootstrap1, (*Acropora clade2*) bootstrap2), *Astreopora* sp1), defined as gene duplication topology (Fig 2c).

We performed comprehensive phylogenomic analysis to confirm IAsα. First, we defined orthogroups as clusters of homologous genes in *Acropora* derived from a single gene in *Astreopora* sp1. Each orthogroup contained at least seven homologous genes, including at least one gene copy in each *Acropora* species and one gene copy in *Astreopora* sp1. We selected 883 orthogroups from 19,760 gene families, and reconstructed the phylogeny of 883 orthogroups using both Maximum likelihood (ML) and Bayesian methods. We found that the phylogeny of 205 orthogroups was consistent with gene duplication topology supporting IAsα. We further defined the 205 orthogroups as core-orthogroups (Supplementary Table 3, See Additional file 1).

In particular, we found differential gene loss, retention, and duplication in *Acropora* lineages. For instance, the phylogeny of orthogroup 1370 (alpha-protein kinase 1-like) showed gene retention in *A. subgrabla, A. digitifera*, and *A. echinata*, gene loss in *A. tenuis*, and an extra gene duplication in *A. subgrabla*. This implies that diversification of duplicated genes may contribute to species complexity and evolutionary innovation in *Acropora* [22] (Fig 2c).

In order to estimate the split time of the two *Acropora* clades that could be regarded as the timing of IAsα, we selected 154 high-quality core-orthogroups, with both bootstrap values in both *Acropora* clades > 70 in ML phylogeny, to reconstruct a time-calibrated phylogeny from the 205 core-orthogroups using BEAST2 [23]. However, we found that it is difficult for the parameters in MCMC to converge in 70 core-orthogroups, and we successfully dated the phylogenetic trees of only 135 high-quality core-orthogroups (See Additional file 1). Next, we estimated the distribution of inferred node ages between the two *Acropora* clades and the peak value was estimated as 30.78 My (95% confidence interval: 27.86-34.77 My), indicating that IAsα occurred at 30.78 My (Fig 2d). This result strongly supports the timing of the IAsα estimated using the dS-based method.

Intergenomic co-linearity is often used to directly identify ancient WGD and to reconstruct ancestral karyotypes in vertebrates [48,53,55]. We performed intergenomic co-linearity and synteny analysis between *Astreopora* sp1 and *A. tenuis* to support IAsα. First, we found great co-linearity between *Astreopora* sp1 and *A. tenuis*. Second, more duplicated scaffold segments were observed in *A. tenuis* than in *Astreopora* sp1 (Supplementary Fig 6) and we also found that some duplicated regions were located in different scaffolds in *A. tenuis* (Supplementary Fig 6).

In summary, we clearly established IAsα using the dS-based method, phylogenomic and synteny analyses. Moreover, we suggest that IAsα probably occurred between 27.86 and 35.7 Mya (Fig 1b).

### The fate of duplicated genes originating from IAsα

Duplicated genes provide substrates for diversification and evolutionary novelty, and most of them are regulators of complex gene networks in vertebrates and plants [23,48,56]. We examined gene ontology (GO) for all genes among the 154 high-quality core-orthogroups to investigate their roles in IAsα and found that their molecular functions have been enriched in specific categories; transporter, catalytic, binding, and receptor activity, most of which are involved in gene regulation (Table 1).

**Table 1.** Gene ontology clusters of 154 high-quality core-orthogroups.

| Annotation cluser | P_Value |
| --- | --- |
| Transmembrane | 1.90E-06 |
| Death domain | 3.10E-05 |
| G-protein coupled receptor | 1.20E-04 |
| VIT domain | 3.30E-03 |
| Protein kinase-like domain | 1.90E-02 |

Further, we identified some duplicated genes under subfunctionalization and neofunctionalization, possibly contributing to stress responses of corals. dnaJ homolog subfamily B member 11-like (DNAJB) protein was shown to be involved in heat stress responses in marine organisms [57,58]. Orthogroups 1247 (DNAJB) has two main domains (Ras and Dnaj domains) in *Astreopora* sp1 representing the ancient state. Each of the two domains was independently lost in the duplicated genes, resulting in complementary functions of the duplicated genes after IAsα (Fig 3a and Supplementary Figs 7). Excitatory amino acid transporters may be related to symbiotic interactions in *Acropora* [59]. Orthogroups 1244 (excitatory amino acid transporter 1-like) was predicted as a six transmembrane protein, and a high number of mutations have accumulated in both untransmembrane and transmembrane regions, suggesting that new functions would be generated (Fig 3b and Supplementary Figs 8). These examples suggest that IAsα participates in both stress responses and symbiotic interactions in *Acropora*. Moreover, these results accord with previous patterns of the fate of duplicated genes in vertebrates and plants [17,19,23,48], indicating that the IAsα possibly contributes to the species complexity and diversification in *Acropora*.

**Figure 3.**
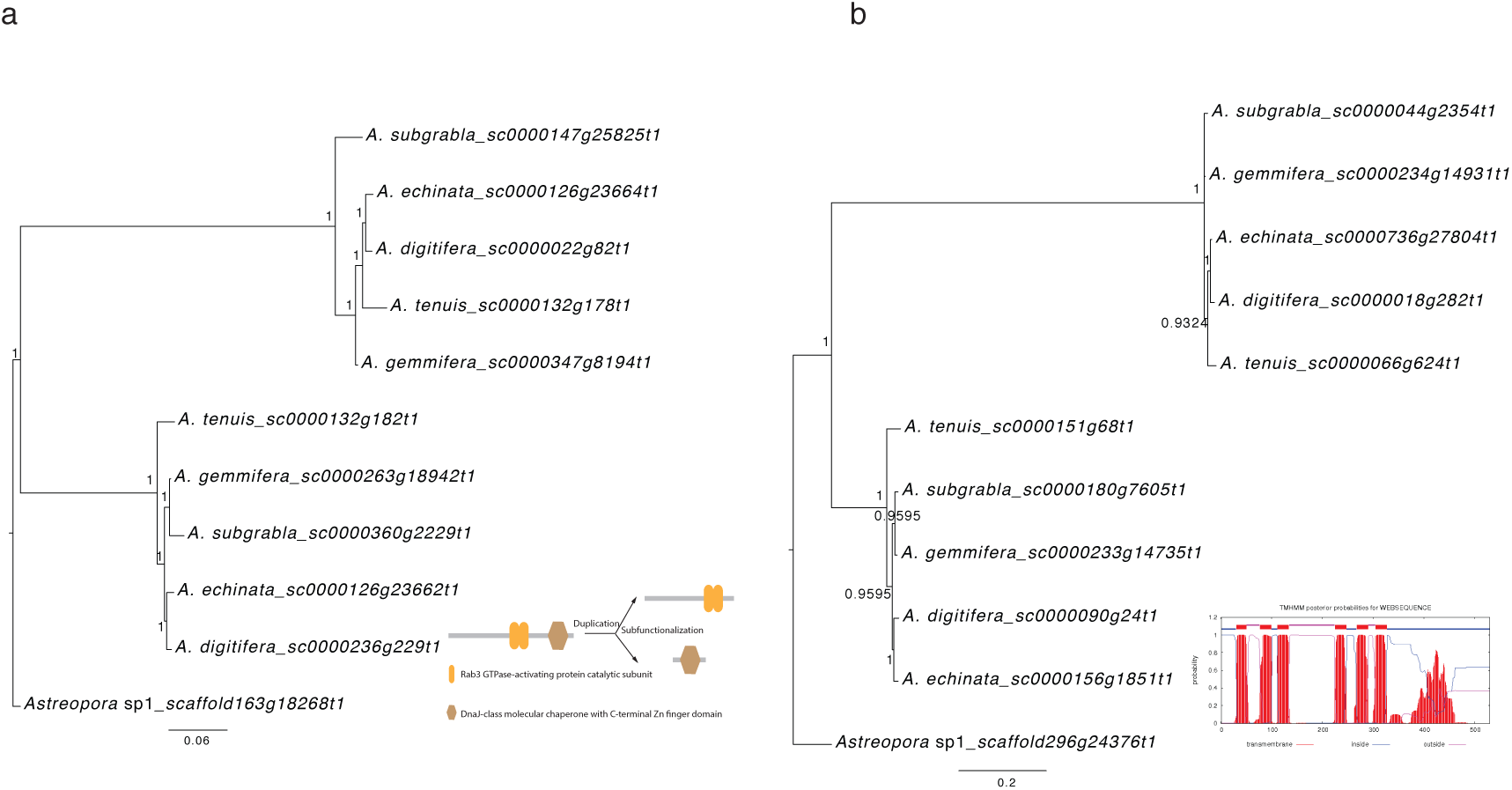
Phylogenetic trees show duplicated genes under subfunctionalization or neofunctionalization. (a). The phylogeny of orthogroup 1247 (dnaJ homolog subfamily B member 11-like) reconstructed with MrBayes shows duplicated genes under subfunctionalization. Bayesian posterior probabilities are shown at each node. The bottom right panel shows that two domains are in *Astreopora* sp1, but each domain was independently lost in duplicated genes under subfunctionalization in orthogroups 1247. (b). The phylogeny of orthogroup 1244 (excitatory amino acid transporter 1-like) reconstructed with MrBayes shows duplicated genes under neofunctionalization. Bayesian posterior probabilities are shown at each node. Six transmembrane helices prediction is shown in the bottom right.

### Gene expression patterns of duplicated genes across five developmental stages in A. digitifera

To better to understand evolution of duplicated genes, gene expression analysis across five developmental stages in *A. digitifera* (blastula, gastrula, postgastrula, planula, and adult polyps) was carried out based on previous transcriptome data [60]. We identified 236 ohnologous pairs in *A. digitifera* from 883 ML phylogeny (for details see Methods) and found that these ohnologous pairs present an interesting gene expression profiling. We divided 236 ohnologous pairs into two clusters based on the pairwise correlation of gene expression during development (high correlation or HC: P<0.05; no correlation or NC: P>=0.05; Pearson’s correlation test); 25% (25/236) ohnologous pairs in HC and 75% (211/236) ohnologous pairs in NC (Fig 4a). Ohnologous pairs in the HC cluster are enriched in protein kinase, while ohnologous pairs in the NC cluster are enriched in membrane transporter and ion binding proteins (Fig 4b). This result indicates that the two clusters of ohnologous pairs potentially evolved into different gene functions. Additionally, we compared dN/dS values in order to investigate selective pressure between HC and NC clusters (Fig 4c), but there is no significant difference between the two clusters (Mann-Whitney-Wilcoxon Test, P = 0.51).

**Figure 4.**
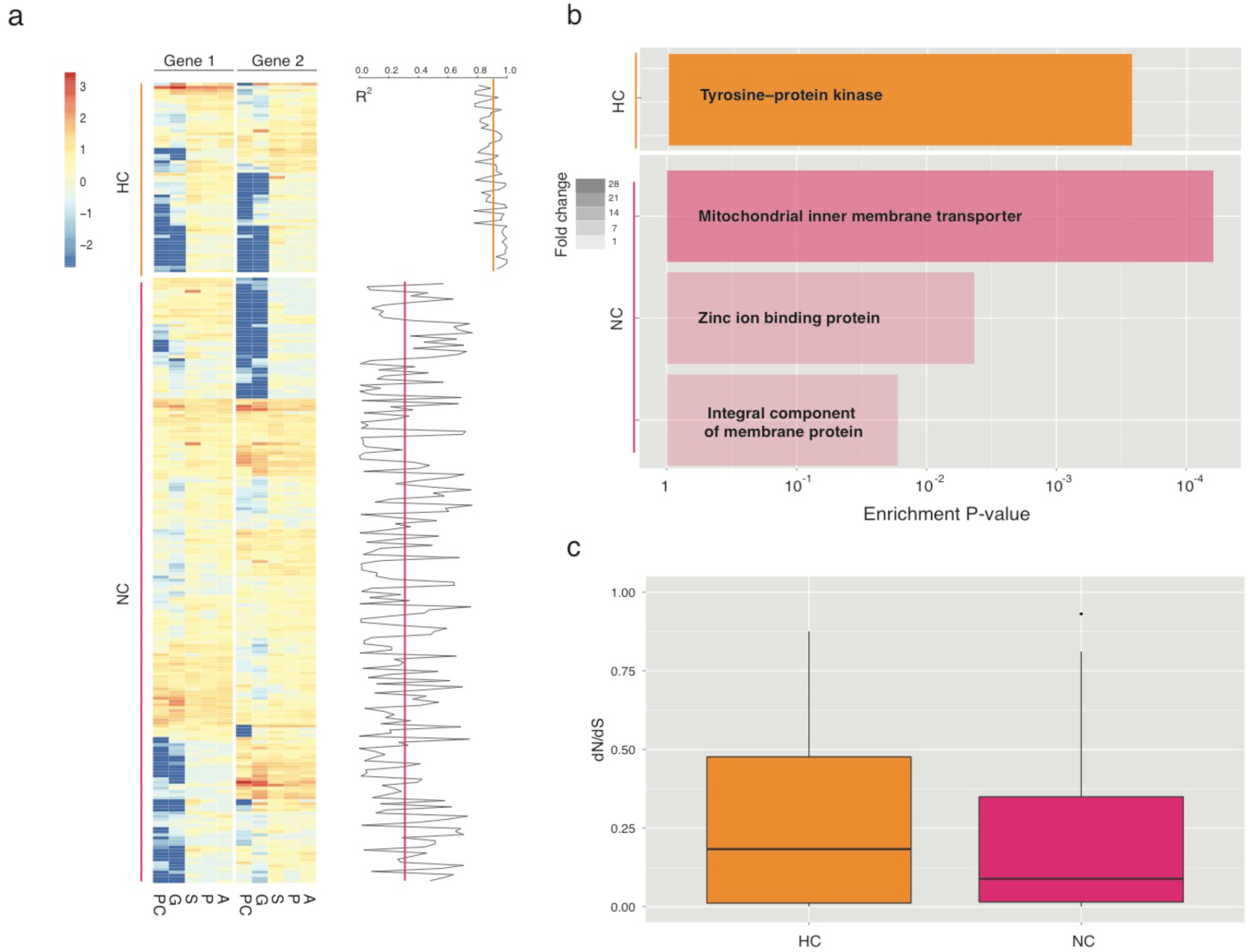
Gene expression profiling reveals evolution of duplicated genes in *A. digitifera*. (a). Gene expression profiling across five developmental stages (blastula: PC, gastrula: G, postgastrula: S, planula: P, and adult polyps: A) in *A. digitifera*. Two clusters of gene expression of ohnologous gene pairs: HC: high correlation, P<0.05; NC: no correlation, P>=0.05 (Pearson’s correlation test). Pearson’s correlation coefficients between two ohnologous gene pairs are presented in the right panel and lines represent average values of correlation coefficients in each cluster. (b) Significant functional enrichments of two clusters of ohnologues gene pairs (P<0.05, Fisher’s exact test) indicate that divergence of gene expression is associated with gene functions. Colors of the bar represent fold change values in enrichments. (c) Boxplot of dN/dS values of ohnologous gene pairs shows no significant difference between the two clusters (P=0.51, Mann-Whitney test).

### Evolution of toxic proteins in Cnidaria

Next, we investigated the role of IAsα in the diversification of toxins in *Acropora*. We identified about two hundred putative toxic proteins in each of the five *Acropora* species, and then we clustered them with putative toxic proteins of *Astreopora* sp1 and other six Cnidarian species (*Hydra magnapapillata, Nematostella vectensis, Montastraea cavernosa, Porites australiensis, Porites astreoides, and Porites lobata*) into 24 gene families (Supplementary Table 4, for details see Methods). Based on reconstruction of the gene family phylogeny, which contains at least 15 genes (See Additional file 1), we found that toxic proteins have undergone widespread gene duplications in Cnidaria, and most of gene duplications occurred in individual species lineages, except for *Acropora* (Fig 5 and Supplementary Figs 9-16). Interestingly, gene duplications occurred in the most recent common ancestor of *Acropora* in 9 of 15 gene families, potentially caused by WGD (IAsα). For example, in gene family-1 (Coagulation factor X), each species contains around 50 genes, except *H. magnapapillata*, and gene duplications occurred frequently in individual species lineages: *Astreopora* sp1, *M. cavernosa, N. vectensis*, and *P. australiensis*. However, five gene duplications occurred in the most recent common ancestor of *Acropora* by WGD (Fig 5). These results indicated that IAsα contributed to the diversification of proteinaceous toxins in *Acropora*.

**Figure 5.**
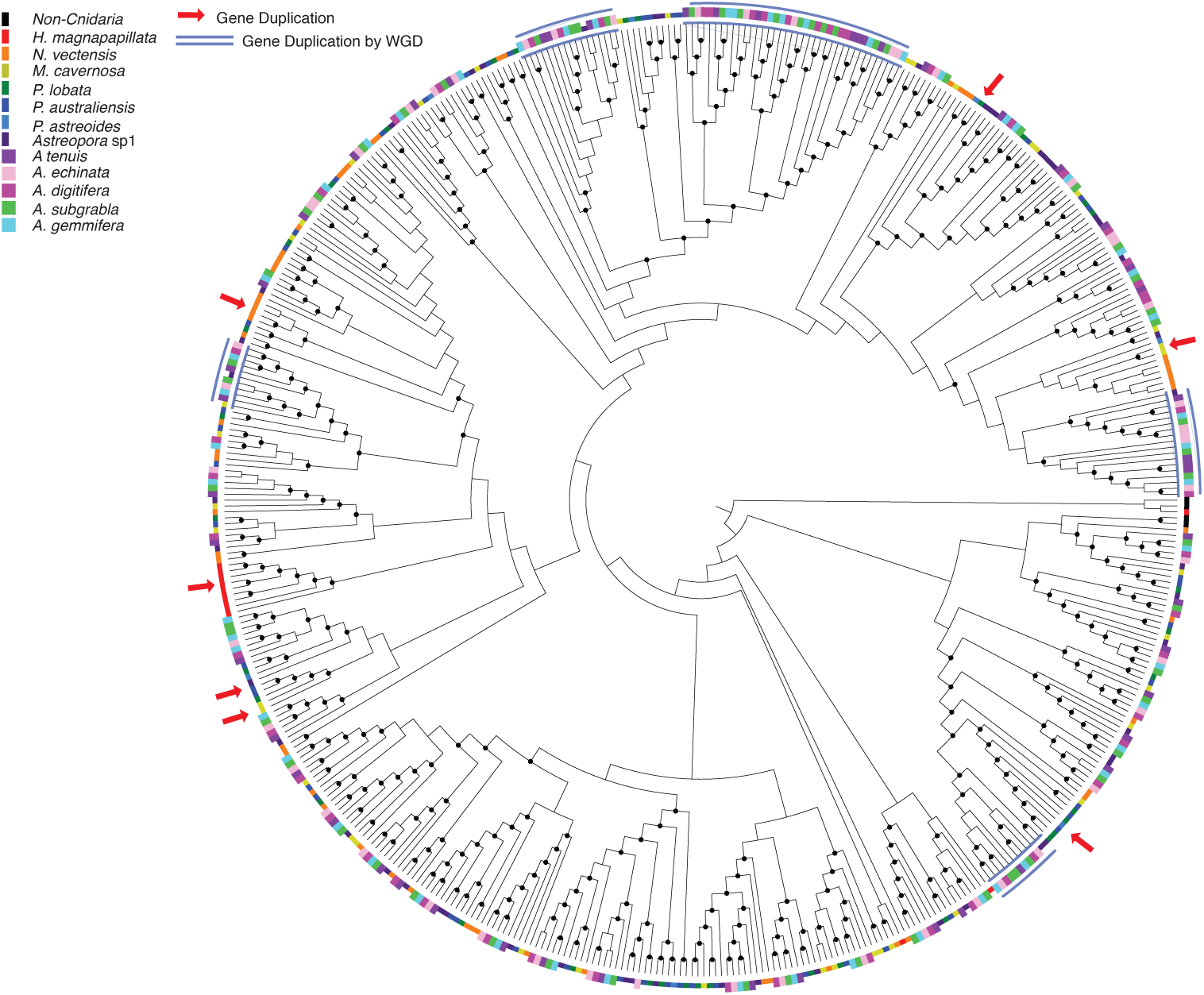
Diversification of toxic proteins via gene duplications in Cnidaria. Phylogenetic analysis of Coagulation factor X in 12 Cnidarian species shows widely gene duplications. Gene duplication occurred in individual species lineages (red arrows) and gene duplications by WGD in *Acropora* are indicated with blue arches. Outer color strips represent 12 Cnidarian species and black strip represents Non-Cnidarian species. Bootstrap values greater than 50 are shown with black dots at nodes.

## Discussion

Ancient WGD is considered as a significant evolutionary factor in the origin and diversification of evolutionary lineages [17,18,28,54], but much work remains to definitively identify WGD and to understand its consequences in different evolutionary lineages. Staghorn corals of the genus *Acropora*, which constitute the foundation of modern coral reef ecosystems, are hypothesized to have originated through polyploidization [2,11,15]. However, there is no genetic evidence to support this assertion. To that end, we analyzed genomes of one *Astreopora* and five *Acropora* species to address the possibility of WGD in *Acropora* and the functional fate of duplicated genes from that event.

To the best of our knowledge, this is the first study to report genomic-scale evidence of WGD in corals (IAsα). We find that large numbers of ohnologs are retained in *Acropora* species and hundreds of gene families display phylogenetic duplication topology among the five *Acropora* species, meanwhile, our synteny analysis between *Astreopora*. sp1 and *A. tenuis* directly supports IAsα. However, reconstruction of the ancestral karyotype will necessitate genomes assembled to the chromosome level to fully understanding gene fractionation and chromosome arrangements in *Acropora* under IAsα [27,61].

Ancient WGD is usually inferred using the dS-based method, but artificial signals in dS distributions have been reported in previous studies, because of dS saturation (dS value > 1) or because of using poorly annotated genomes [24,52,62]. There is an extra peak in the dS distribution of anchor gene pairs in *A. digitifera* and *A. tenuis*. One possible explanation is that the extra peak is artifactitious because few anchor gene pairs were used in the analysis. However, this could also indicate a second WGD event in *Acropora*. We found few orthogroups with topologies that fit the two proposed WGDs events (Supplementary Fig 17). If a second WGD event occurred, the reason that the second WGD signal appeared among anchor gene pairs rather than among paralogous gene pairs may be that the paralogs generated by the second WGD have been largely lost; thus, few of them are only retained in conserved order. Fortunately, a new maximum likelihood phylogeny modeling approach was recently developed to overcome difficulties of the dS-based method [24,62]. We used it to test whether a second WGD occurred in *Acropora*. The result showed that one WGD event is the best model in *Acropora* and it occurred 30.69 to 34.69 Mya (Supplementary Table 5 and Supplementary Table 6; for details see Methods). Thus, we have supportive genome-scale evidence to support IAsα, but as yet, there is no conclusive evidence to support a second WGD in *Acropora*.

It is crucial to accurately estimate the timing of a WGD event to understand its evolutionary consequences [23,25]. Our study has clearly estimated the timing of IAsα using both phylogenomic analysis and the dS-based method. We suggest that IAsα probably occurred between 27.86 and 35.7 Mya (Fig 1b). Interestingly, species turnover events usually occurred with extinctions [63], and one species turnover event in corals (Oligocene-Miocene transition: OMT) was suggested to have occurred from 15.97 to 33.7 Mya [8]. The timing of IAsα may correspond to a massive extinction of corals created by OMT. This finding supports the hypothesis that WGD may enable organisms to escape extinction during drastic environmental changes [17] (Fig 1b).

The occurrence of IAsα raises the question of what impact it may have had upon coral evolution [15,32]. We performed GO analysis on duplicated genes and examined several duplicated gene families, showing that duplicated genes following IAsα indeed provided raw genetic material for *Acropora* to diversify and are potentially crucial for stress responses. In particular, toxin diversification in *Acropora* was mainly generated by WGD. In addition, we focused on expression patterns of duplicated genes in *A. digitifera*, showing that expressions of duplicated protein kinases are likely to be correlated during development. A possible explanation may be that protein kinases are probably retained in complex signal transduction pathways via subfunctionalization or dosage effects [22,32]. However, expressions of duplicated membrane proteins are likely uncorrelated probably because these proteins may have developed different functions via neofunctionalization, such as excitatory amino acid transporters (orthogroups 1244). However, there is still much work needed to investigate molecular mechanisms of duplicated genes to examine these hypotheses in the diversification of *Acropora* [64]. For instance, previous gene functional studies have demonstrated that voltage-gated sodium channel gene paralogs, duplicated in teleosts, contributed to the acquisition of new electric organs via neofunctionalization in both mormyroid and gymnotiform electric fishes [65,66].

Our previous study proposed that adaptive radiation in *Acropora* was probably driven by introgression (Mao et al., in revision); thus, *Acropora* is the first invertebrates lineage reported to have undergone both WGD and introgression. Meanwhile, both introgression and WGD have also been reported in cichlid fish lineages [67], a famous model for adaptive radiation in vertebrates [67,68]. Both WGD and introgression are regarded as significant forces in adaptive radiation of organisms [17,67], but we still do not understand the relationship between WGD and introgression in adaptive radiations [69].

In conclusion, this study identified an ancient WGD shared by *Acropora* species (IAsα) that not only provides new insights into the evolution of reef-building corals, but also expands a new model of WGD in animals.

## Methods

### Species information, genomic data and gene families cluster

Data can be accessed at: http://marinegenomics.oist.jp and http://comparative.reefgenomics.org/datasets.html. The *Acropora* species information in this study will be described in the paper (Mao et al., in revision). Information about *Astreopora* sp1 was described previously [7] and transcriptome data across five development stages was descried previously [60]. Protein sequences of the six species were combined together to perform all-against-all BLASTP approach to find all orthologs and paralogs among six species. Then, OrthoMCL was used with default settings to cluster homologs into 19,760 gene families according to sequence similarity [70].

### Single-copy orthologs and reconstruction of a calibrated phylogenomic tree

A custom python script was used to select 3,461 single-copy orthologs with only one gene copy in each species. For each sequence alignments of single-copy orthologs, coding sequences were aligned with MAFFT [71] as described previously. Then, the concatenated sequences of 3,461 single-copy orthologs were used to reconstruct the phylogenomic tree (species tree) with BEAST2 [72]. First, we partitioned the concatenated coding sequences by codon position. Molecular clock and trees, except substitution model, were linked together. Then, divergence time was estimated using the HKY substitution model, relaxed lognormal clock model, and calibrated Yule prior with the divergence time estimated in our previous study (Mao et al., in submission). We ran BEAST2 three times independently, 50 million Markov chain Monte Carlo (MCMC) generations for each run, then we used Tracer to check the log files and we found that ESS of each of parameters exceeded 200.

### Orthogroup selection and detection of a WGD event with dS analysis

#### (a) dS distributions of paralogous gene pairs

Paralogous gene pairs of each species were identified by all-against-all BLASTP approach and then OrthoMCL was used to cluster paralogs to gene families for each species [70]. Gene families with fewer than 20 genes were used to calculate dS values. Each gene pair within a given gene family was aligned with MAFFT [71] and aligned sequences were used to calculate dS values with PAML [73]. The dS distribution of each species was plotted with bins=0.02 in R [74].

#### (b) dS distributions of anchor gene pairs

First, we used MCScanX with default settings (except for match_size=3) to find anchor gene pairs based on synteny information for each species [75]. Each anchor gene pair was aligned with MAFFT [71] and aligned sequences were used to calculate dS values with PAML [73]. The dS distribution of each species was plotted with bins=0.02 in R [74].

#### (c) dS distributions of orthologous gene pairs

First, we used MCScanX with default settings (except for match_size=3) to find orthologous gene pairs based on synteny information between *Astreopora* sp1 and *A. tenuis*, and between *A. tenuis* and *A. digitifera* [75]. Each orthologous gene pair was aligned with MAFFT [71] and aligned sequences were used to calculate dS values with PAML [73]. dS distributions of all species were plotted with bins=0.02 in R [74].

### Detection of a WGD event using phylogenetic analysis

A custom python script was used to select the 883 gene families, including at least one gene copy in *Astreopora*, one gene copy in each of the five species and at least two ohnologs in one of five *Acropora* species, as orthogroups. Ohnologs are defined as paralogs originating from WGD.

For each of the 883 gene tree reconstructions, we used MAFFT [71] to align amino acid sequences of each single-copy ortholog. We aligned coding sequences with TranslatorX [76] based on amino acid alignments and we excluded the single-copy orthologous genes containing ambiguous ‘N’. PartitionFinder [77] was used to find the best substitution model for RAxML (Version 8.2.2) [78] and MrBayes (Version 3.2.3) [79]

Then, 205 orthogroups, for which phylogeny matched the duplication topology (*Astreaopora*, (*Acropora, Acropora*)), were selected as core-orthogroups by eyes. The 154 high quality core-orthogroups, for which clades’ bootstrap values in ML phylogeny exceeded 70, were used to perform molecular dating with BEAST2 based on the calibrated phylogenomic tree. Molecular clock and trees, except substitution model, were linked together. Then, divergence time was estimated using the HKY substitution model, relaxed lognormal clock model, and calibrated Yule prior with the divergence time from our previous study (Mao et al., in submission). We ran BEAST2 three times independently, 30 million Markov chain Monte Carlo (MCMC) generations for each run. Then we used Tracer to check the log files. 135 time-calibrated phylogeny with ESS values exceeded 200 were carried out by BEAST2 [72].

### Estimating peak values in dS distributions and inferred node ages’ distribution with KDE toolbox

Each distribution was estimated using KDE toolbox in MATLAB, as described previously [25,80].

#### (a). Estimating peak values in distributions

To estimate the age of WGD within dS distributions, we assumed the peak value in orthologous gene pair dS distributions as the split time between two species: the split time between *Astreopora* sp1 and *A. tenuis* is 53.6 My, whereas that the split time between *A. tenuis* and *A. digitifera* is 14.69 My. Before we used the *kde(*) function in KDE toolbox, we first truncated dS distributions to avoid estimation bias due to extreme values: the dS distribution of orthologous gene pairs between *Astreopora* sp1 and *A. tenuis* was truncated with a range from −1 to1 while the dS distribution of orthologous gene pairs between *A. tenuis* and *A. digitifera* was truncated with a range from −5 to −2. Then, we used the *kde(*) function in KDE toolbox to estimate the peak values of these two dS distributions as −0.314 and −3.4596, respectively. Moreover, the distribution of *Acropora* paralogous gene pairs was truncated with a range from −4 to 0 and we estimated the peak value of this distribution as −1.8165. We also used bootstrapping to estimate 95% confidence intervals (CIs) of *Acropora* paralogous gene pairs distribution as −1.7606 to −2.1261 (31.18 to 35.71 My). For bootstrapping, we generated 100 bootstrap samples for each distribution by sampling with replacement from the original data distribution (49,002 samples in the original distribution) with the *sample(*) function. We estimated maximum peak values for each 100 bootstrap samples. Then we sorted maximum peak values and values of 6^th^ and 95^th^ rank were used to define the 95% CI.

#### (b). Estimating peak values in distributions of inferred node age

To estimate the age of WGD in the distribution of inferred node ages, we used the *kde(*) function in KDE toolbox to estimate the peak value as 30.78 My, and we used bootstrapping to estimate the 95% CIs as 27.86 to 34.77 My. For bootstrapping, we generated 100 bootstrap samples from the distribution by sampling with replacement from the original data distribution (135 samples in the original distribution) with the *sample(*) function. We estimated maximum peak values for each of 100 bootstrap samples. Then, we sorted maximum peak values and values of 6^th^ and 95^th^ rank defined the 95% CI.

### Maximum likelihood approach to detect WGD with gene family count data

First, we filtered gene family cluster data generated by OrthoMCL described above [70]. The gene family, including only one *Astreopora* sp1 gene and at least one gene in each of the five *Acropora* species, was counted. Then, we used the WGDgc package in R to estimate log likelihood for parameters (0, 1, 2, 3) of WGD event(s) with setting (dirac=1,conditioning=“twoOrMore”) [62]. Then, we performed likelihood ratio test (pchisq(2*(Likelihood_1-Likelihood_2), df=1, lower.tail=FALSE)) to find the best model and found that one WGD event was the best model to fit the gene family count data. We estimated the age of WGD on 4 My intervals between 18.69 and 38.69 My under a one WGD event model. The lowest log likelihood was shown at the age of WGD: 30.69 and 34.69 My.

### Gene expression profiling analysis and dN/dS calculation

We selected 236 gene pairs of *A. digitifera* (ohnologous gene pairs) from 831 orthogroups. We BLASTed these ohnologous gene pairs against the gene expression data across five developmental stages [60] and these data were normalized for each developmental stage. Correlations between two ohnologous genes were performed using Pearson’s correlation in R [74]. Hierarchical clustering was performed using Pheatmap for HC cluster genes and NC cluster genes, respectively. Pairwise dN/dS ratios were calculated with PAML using codeml based on the coding sequence alignment of ohnologous gene pairs [73]. The dN/dS distribution was plotted with ggplot2 in R and significance tests of differences between dN/dS distributions were evaluated by a Mann-Whitney test in R [74].

### Evolution analysis of toxic proteins in corals

The 55 toxic proteins of *A. digitifera* identified in the previous study were downloaded from http://www.uniprot.org/ as queries. The proteins sequences of *Porites astreoides, Porites australiensis, Porites lobata, Montastraea cavernosa, Hydra magnipapillata* and *Nematostella vectensis* were downloaded from http://comparative.reefgenomics.org/datasets.html, and combined them with protein sequences of six *Acroporid* species to create a search database.

We identified candidates of toxic proteins by BLASTing the 55 toxins against the combined protein sequences with settings: e-value < 1e^−20^ and identity > 30%. Then, we used OrthoMCL to cluster candidates of toxins into 24 gene families and reconstructed their ML gene trees with ExaML and RAxML. Each gene tree was rooted at a branch or clade of query sequences.

### Gene ontology enrichment for duplicated genes of core-orthogroups and protein domains and transmembrane helices prediction

We BLASTed the sequences of 154 high quality core-orthogroups and ohnologous gene pairs in *Acropora* against the UNIPROT database to find best hits. Identical hits in each ohonlogs group were removed and the remaining hits were used to perform gene enrichment in David [81]. We also used InterProScan [82] to predict protein domains and used the TMHMM Server (v. 2.0) [83] to predict transmembrane helices from protein sequences.

## Acknowledgments

This study was supported by OIST and was funded by JSPS grants (No. 17J00557 to YFM). We thank Dr. Douglas E Soltis and Dr. Pamela S Soltis for their comments and insight into this draft manuscript. We thank Dr. Steven D. Aird for editing the manuscript.

## Availability of data and materials

Genome assembly and gene models in this study can be downloaded from http://marinegenomics.oist.jp.

## Authors’ contributions

YFM and NS conceived the study. CS provided genomic data. YFM performed all analysis in this study. YFM and NS wrote the initial manuscript and all authors edited the final manuscript.

## Additional files

Additional file 1: An Excel file with three tables listing 1) the original data of gene family clusters in this study, 2) orthogroup datasets in this study, 3) node ages of core-orthogrouops inferred by BEAST2 in this study, 4) data of gene expression analysis for ohnologous gene pairs, and 5) toxin gene family clusters.

## Supplementary information

### Supplementary Figures

**Figure S1.**
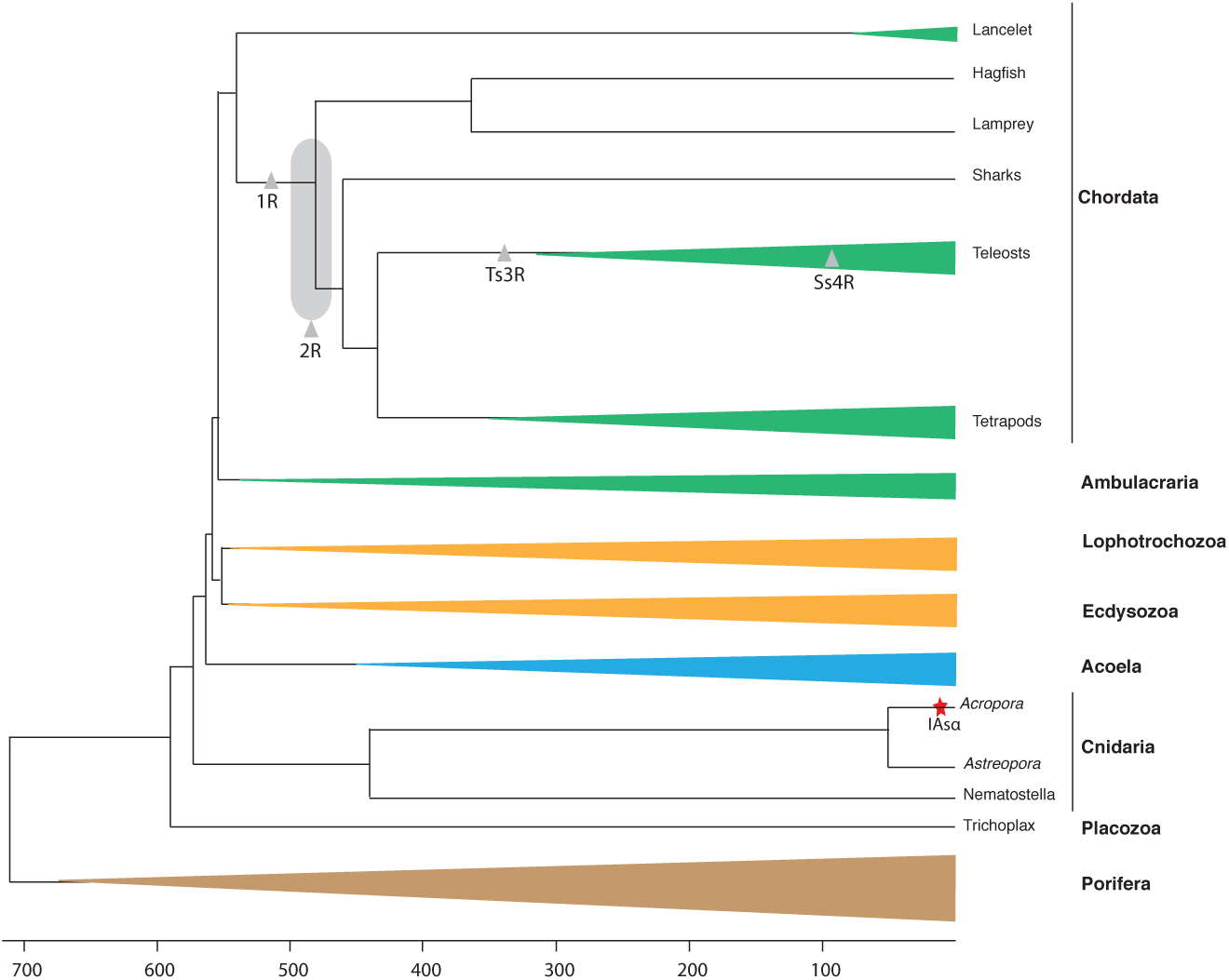
WGD events in evolution of the animal kingdom. The backbone and divergence time of the tree are based on various sources (e.g., Satoh, 2016). The shaded grey oval represents the uncertain position of two rounds of WGD and colored triangles represent the corresponding divergent groups. Grey triangles represent WGD and the red star represents invertebrate WGD specific to *Acropora* (IAsα) reported in this study.

**Figure S2.**
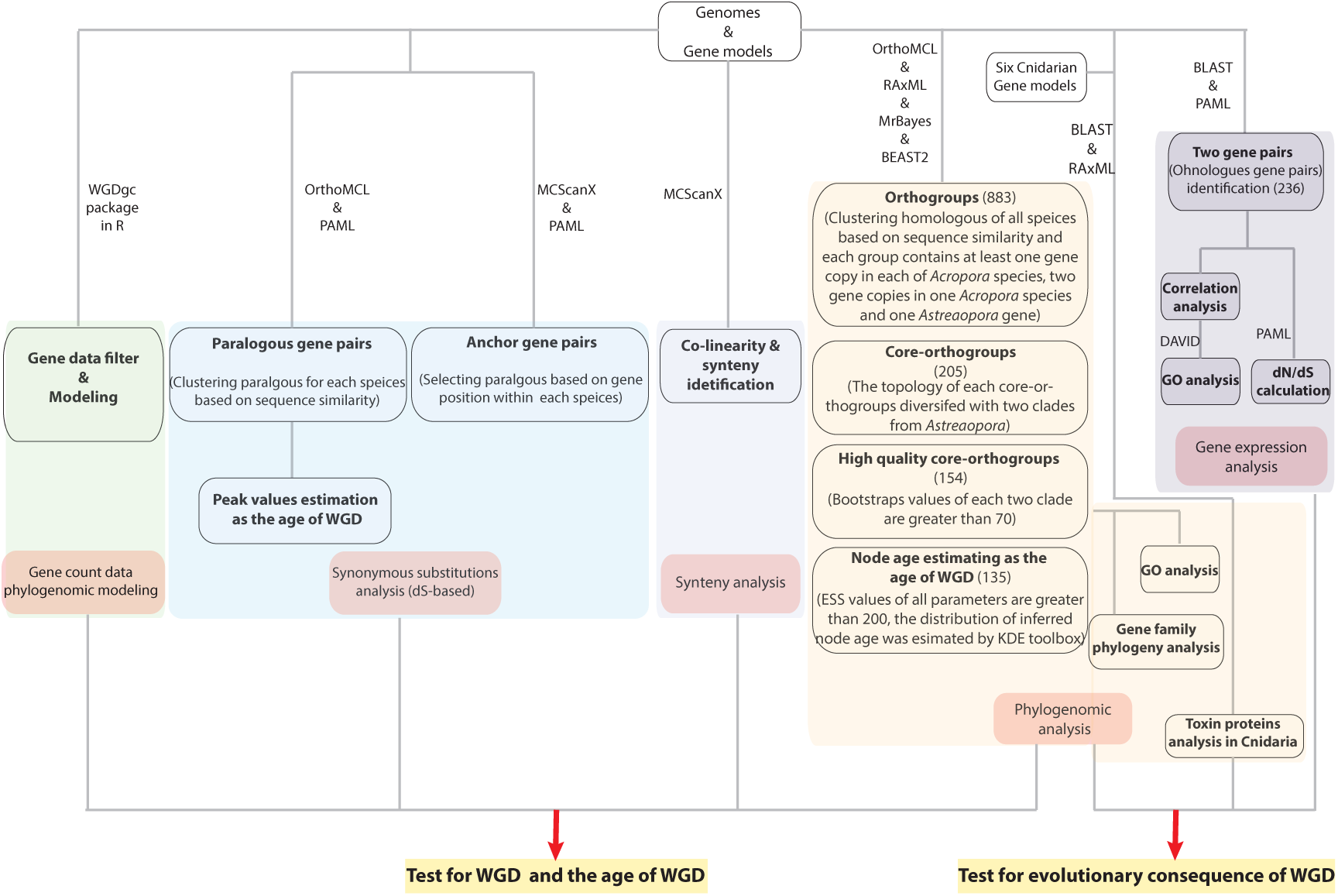
Flowchart of study design. The flowchart illustrates methods used to test the main questions in this study.

**Figure S3.**
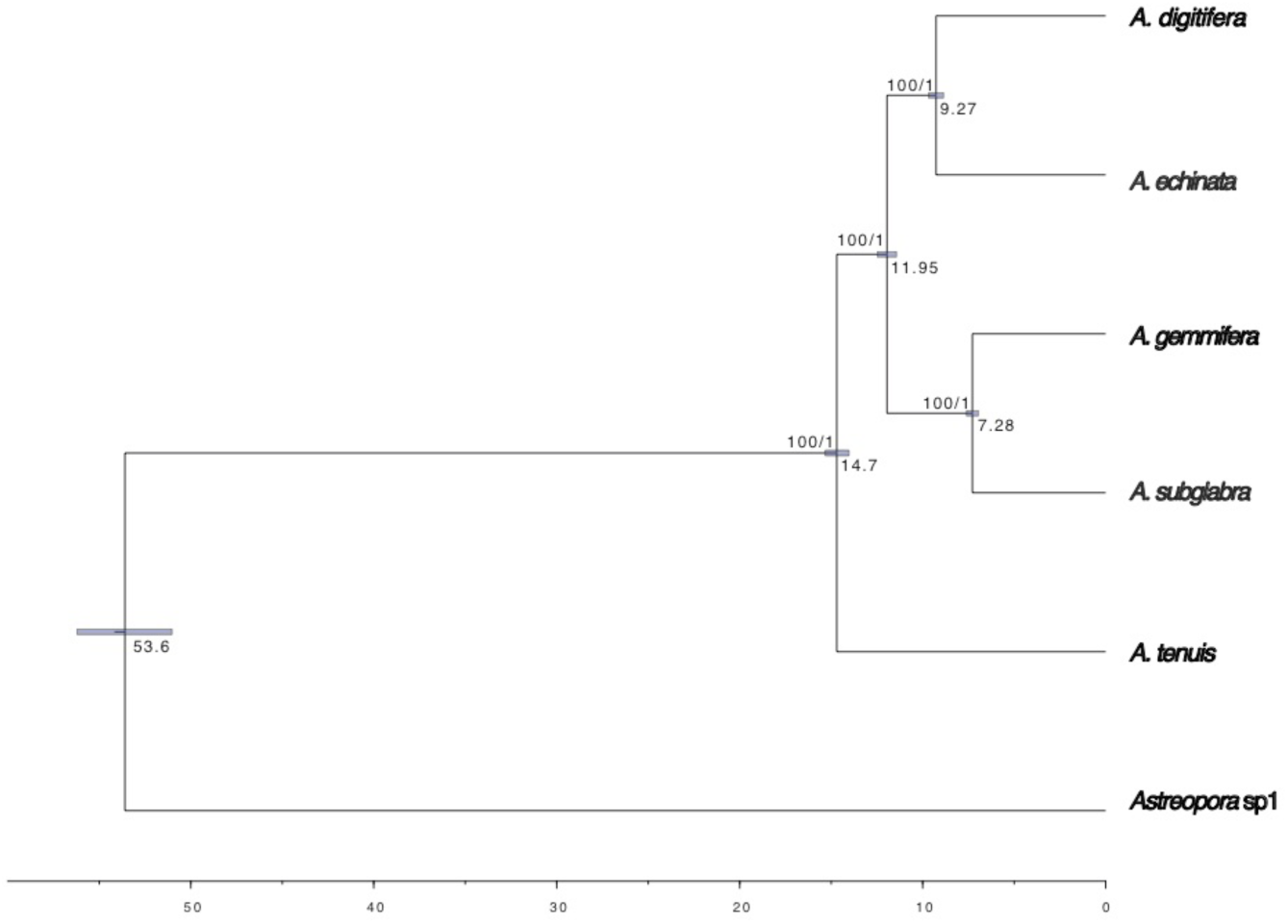
Phylogeny of the Family *Acroporidae*. Time-calibrated phylogenetic tree reconstructed based on fossil calibration and concatenated coding sequences (7,467,066 bp in total) from 3,461 single-copy orthologous genes with Beast2. Branch lengths are scaled to estimated-divergence time. Posterior 95% CIs of node ages are represented with blue horizontal bars as well as ML bootstrap values and Bayesian posterior probabilities are shown at each node.

**Figure S4.**
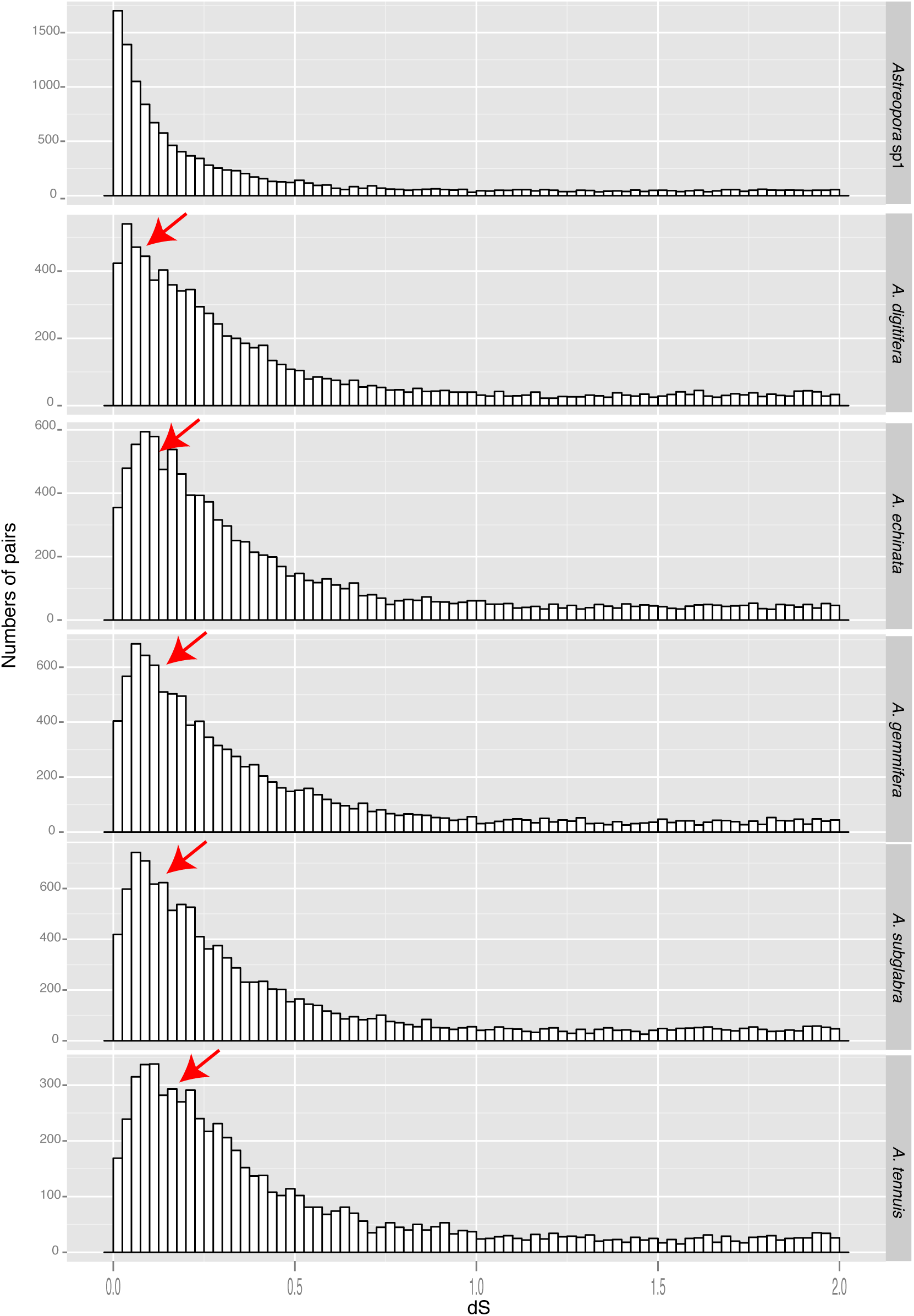
Frequency distribution of dS values for paralogous gene pairs in five *Acropora* and one *Astreopora* species. The distributions of dS values of paralogs, estimating neutral evolutionary divergence since the two paralogs diverged, are plotted with a bin size of 0.02, showing the similar peaks (dS value: 0.1-0.25) in *Acropora* (red arrows).

**Figure S5.**
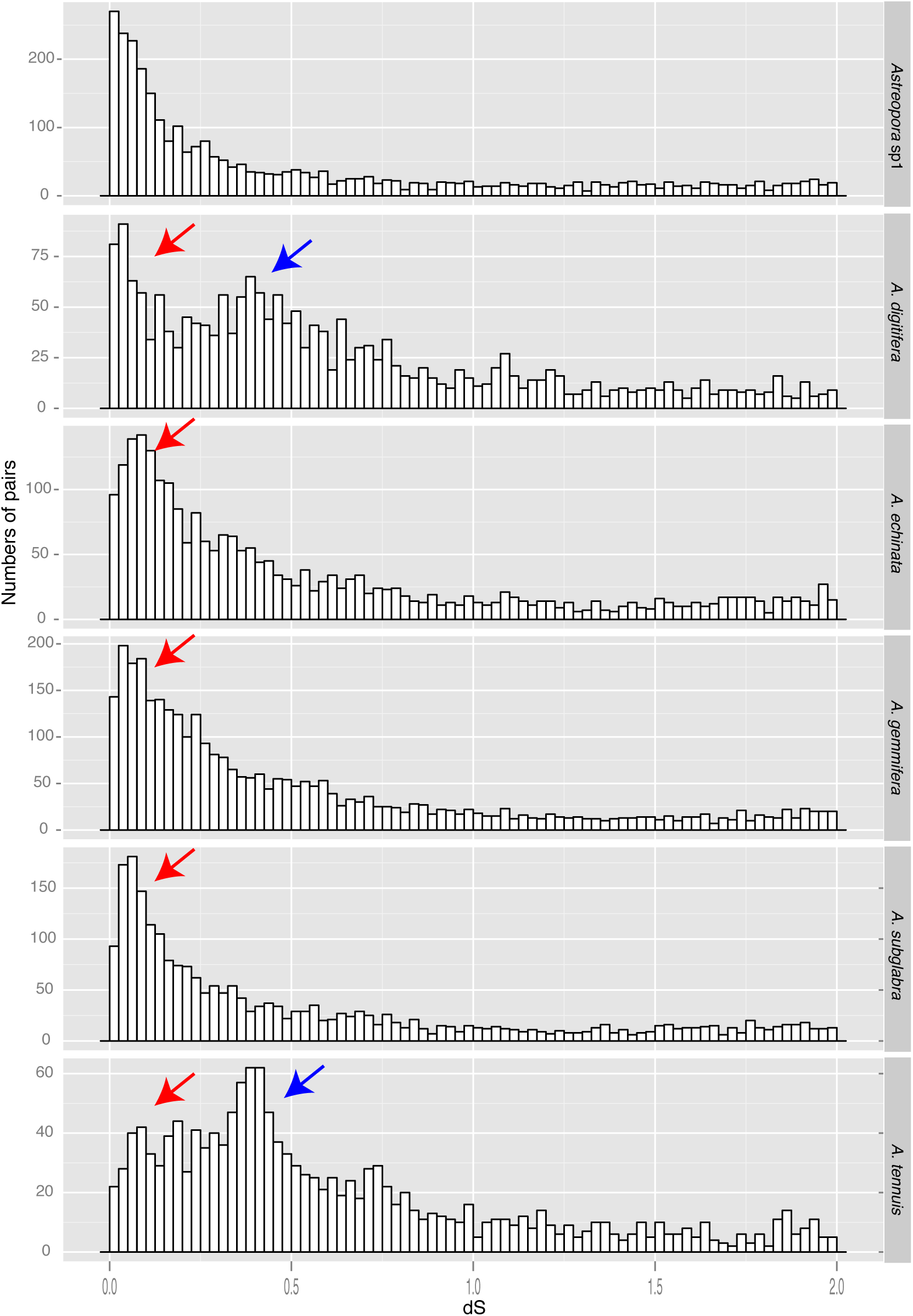
Frequency distribution of dS values for anchor-gene pairs in five *Acropora* and one *Astreopora* species. Distributions of dS values of anchor paralogs, estimating the neutral evolutionary divergence times since the paralogs diverged, are plotted with a bin size of 0.02, showing the similar peaks (dS value: 0.1-0.25, red arrows) in *Acropora* and extra peaks in *A. digitifera* and *A. tenuis* (dS value: 0.25-0.5, blue arrows).

**Figure S6.**
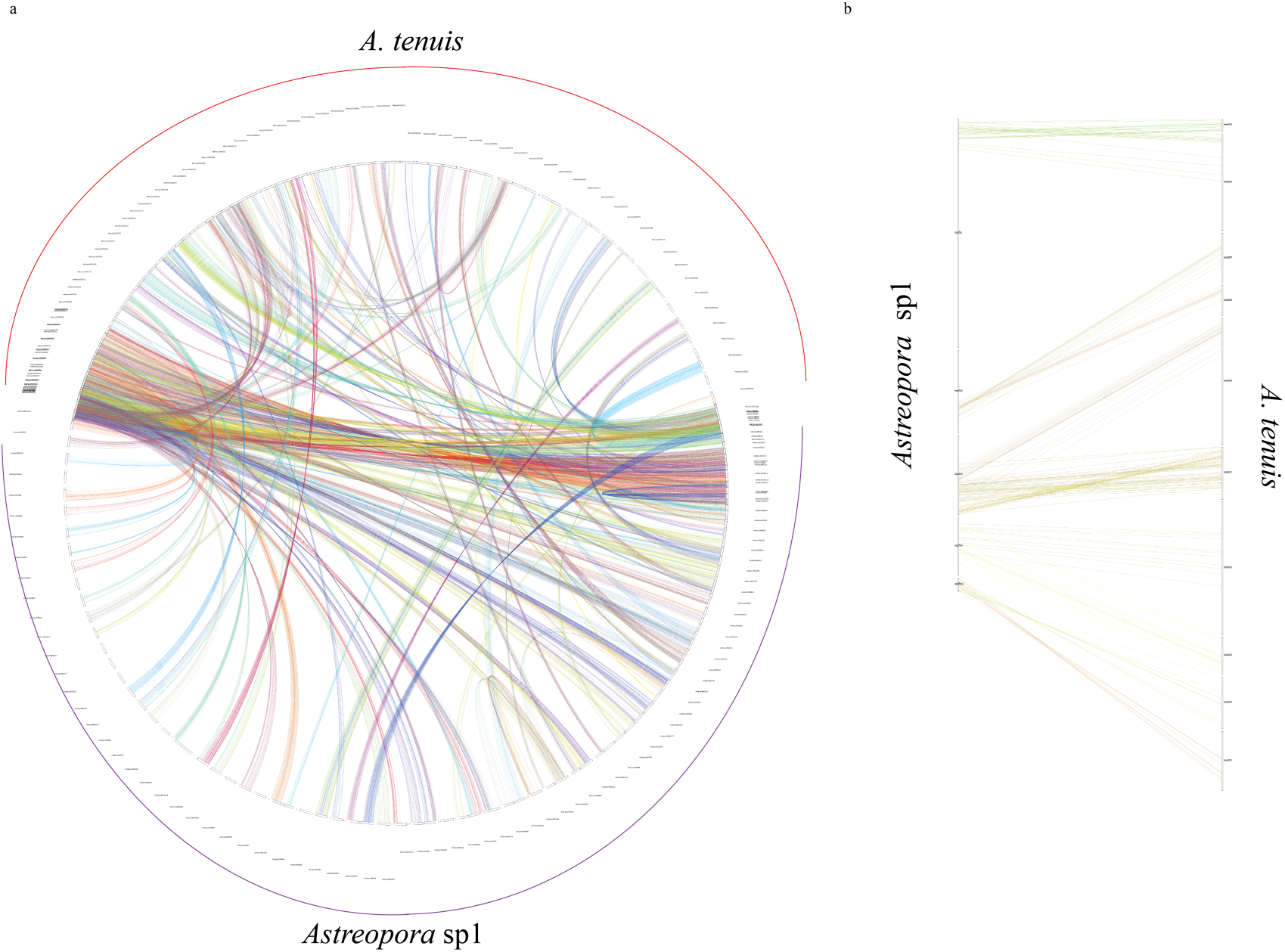
Co-linearity and synteny between *Astreopora* sp1 and *A. tenuis*. Only co-linear segments with at least 8 anchor pairs are shown in the top length 100 scaffolds.

**Figure S7.**
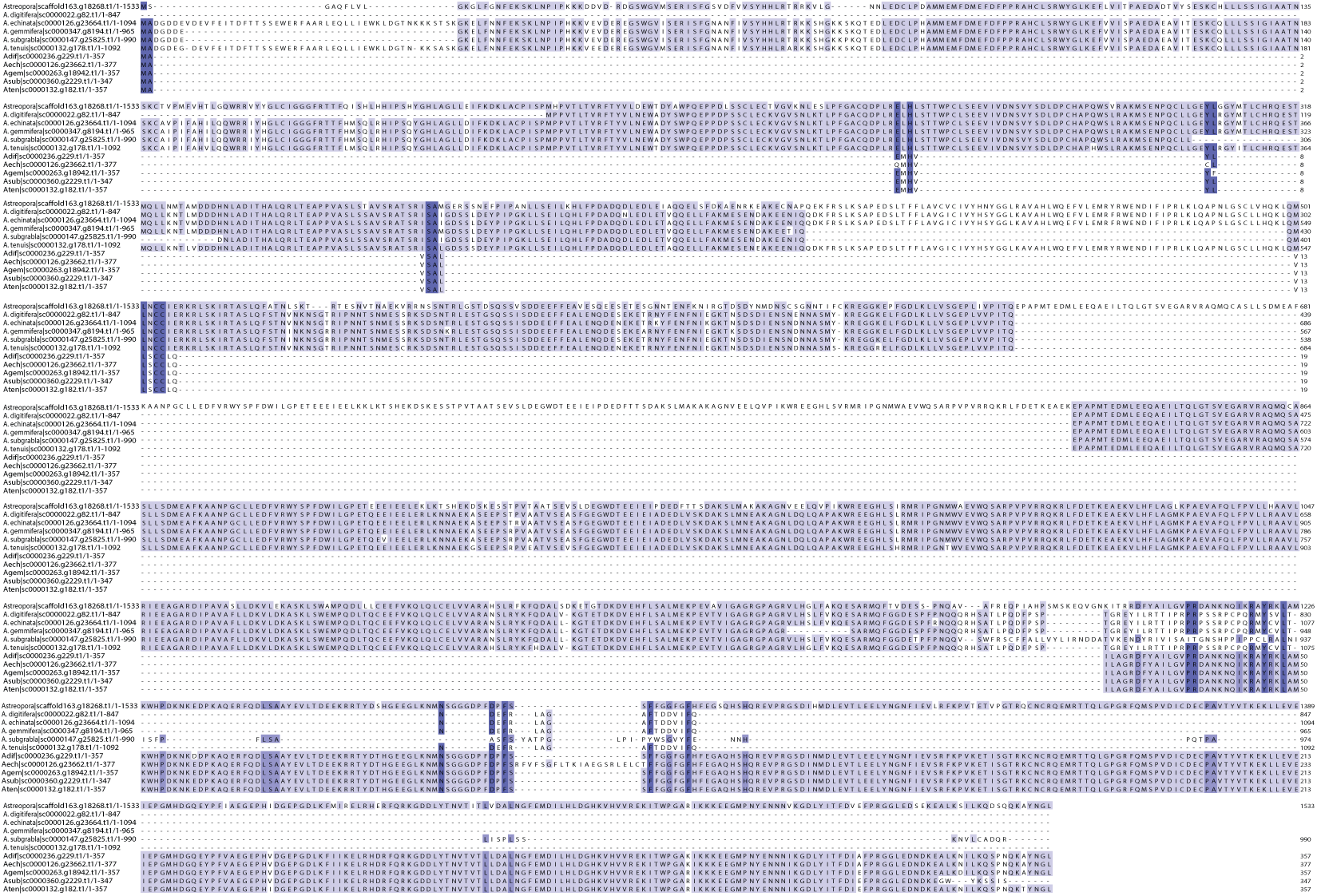
Alignment of orthogroup 1247 (dnaJ homolog subfamily B member 11-like) showing the independent loss of the domain in duplicates.

**Figure S8.**
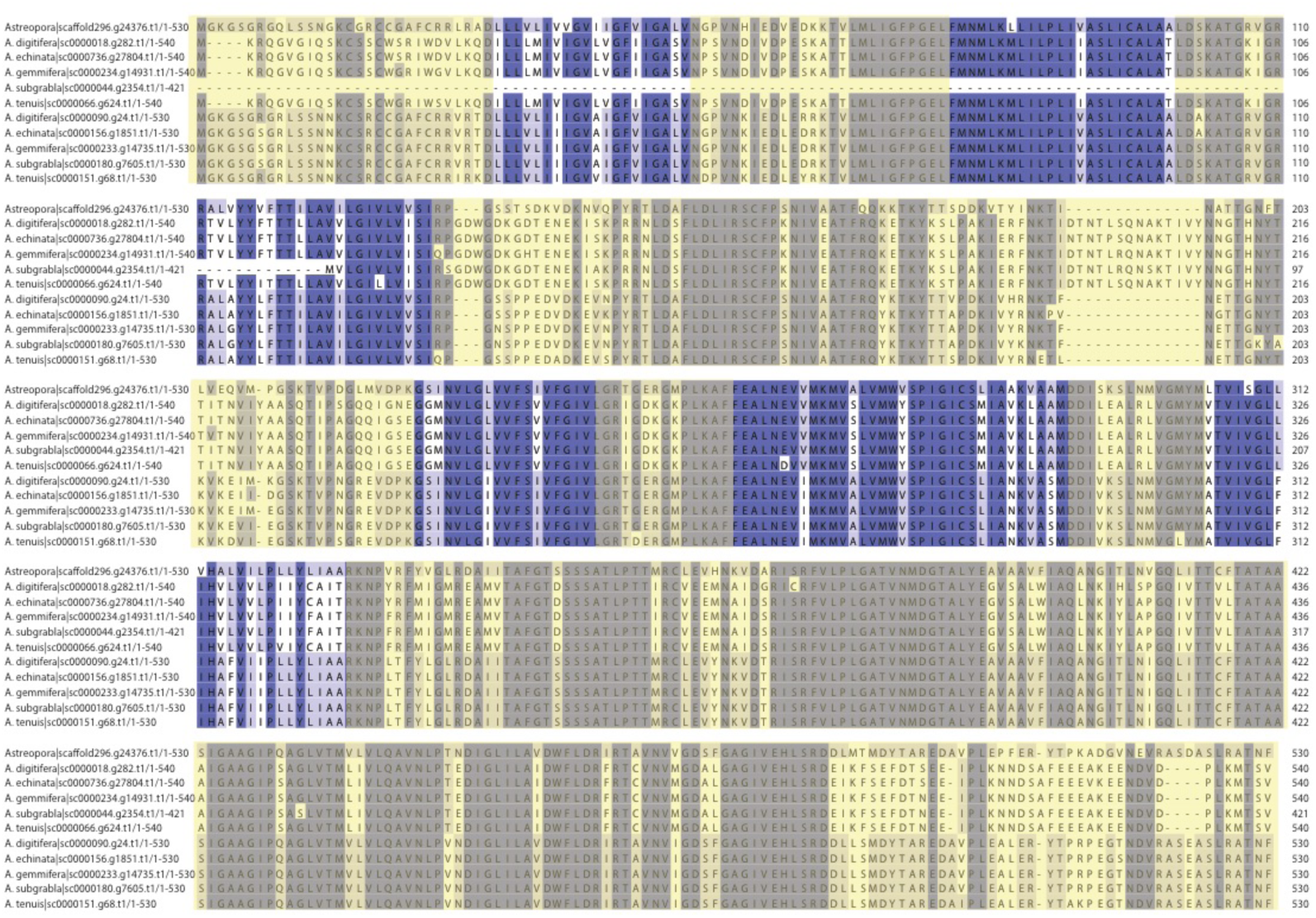
Alignment of orthogroup 1244 (excitatory amino acid transporter 1-like) showing mutations on transmembrane and exposed regions, suggesting that new functions would be generated. Exposed regions are shown in yellow.

**Figure S9.**
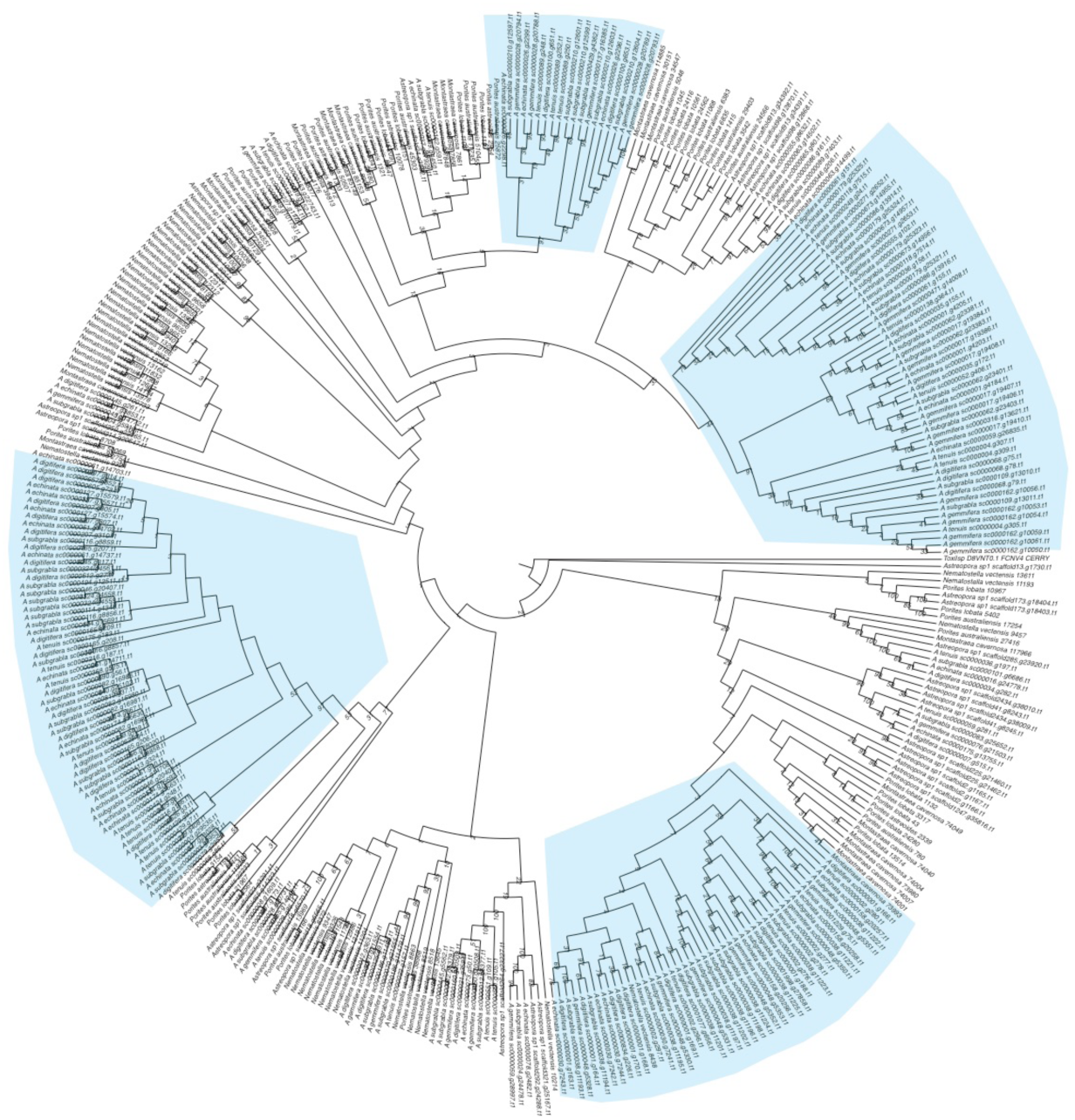
Phylogeny of the toxic protein, Ryncolin-4. The phylogeny was reconstructed using ExaML and bootstrap values are shown at each node. Gene duplications caused by WGD in *Acropora* are shown in cyan.

**Figure S10.**
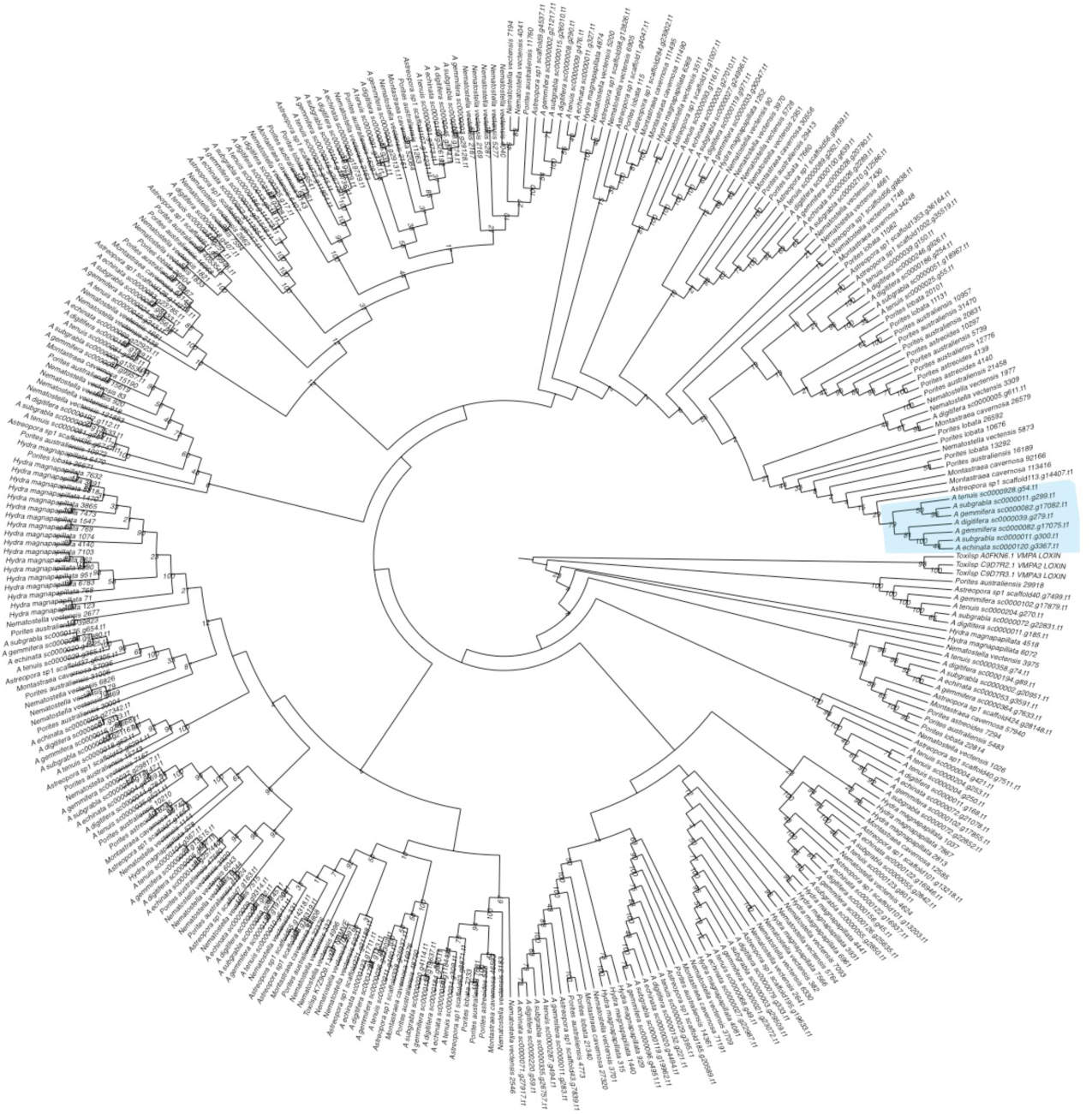
Phylogeny of the toxic Astacin-like metalloprotease. The phylogeny was reconstructed using ExaML and bootstrap values are shown at each node. Gene duplication caused by WGD in *Acropora* is shown in cyan.

**Figure S11.**
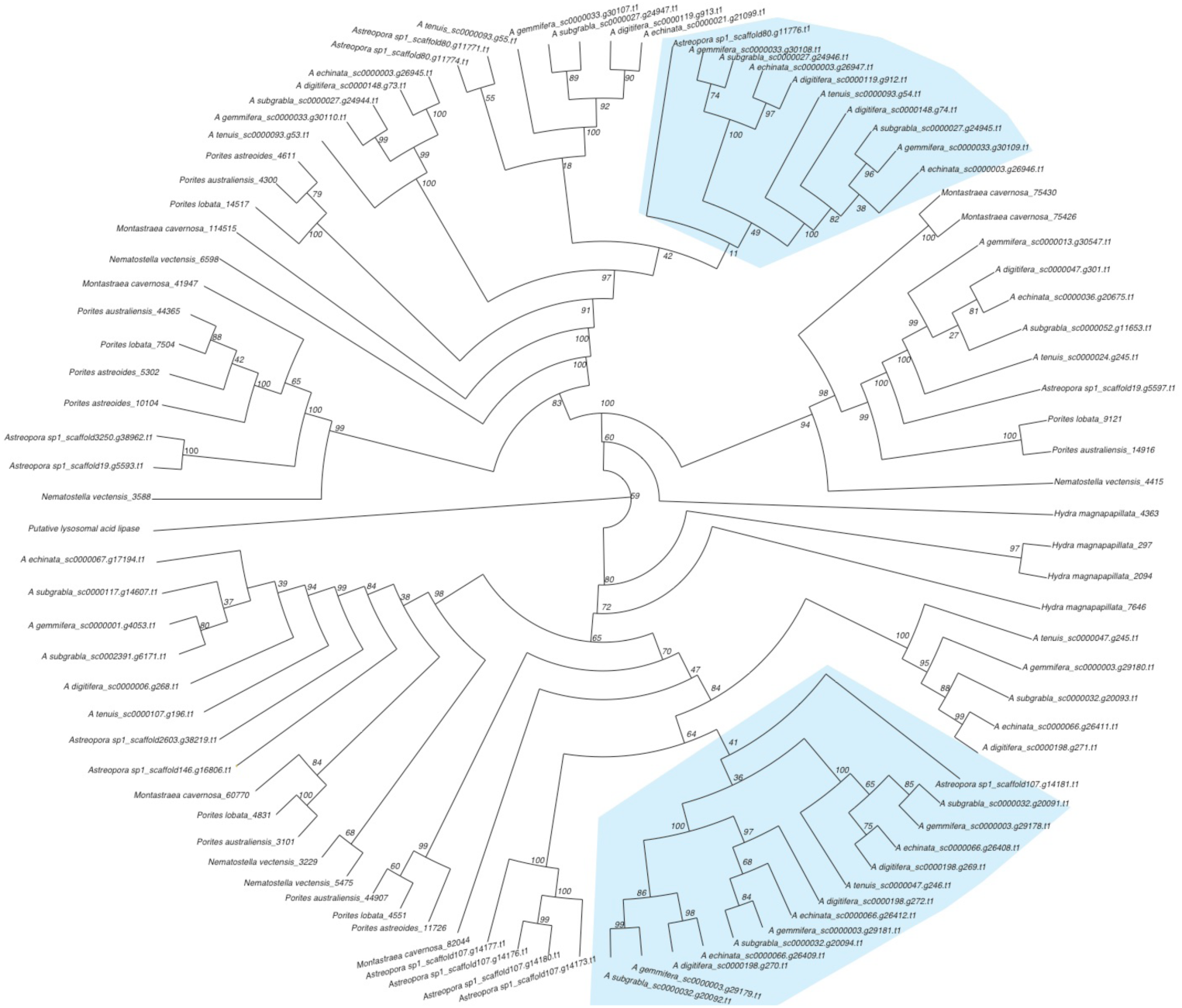
Phylogeny of a toxic protein (Putative lysosomal acid lipase/cholesteryl ester hydrolase). The phylogeny was reconstructed using RAxML and bootstrap values are shown at each node. Gene duplications caused by WGD in *Acropora* are shown in cyan shadows.

**Figure S12.**
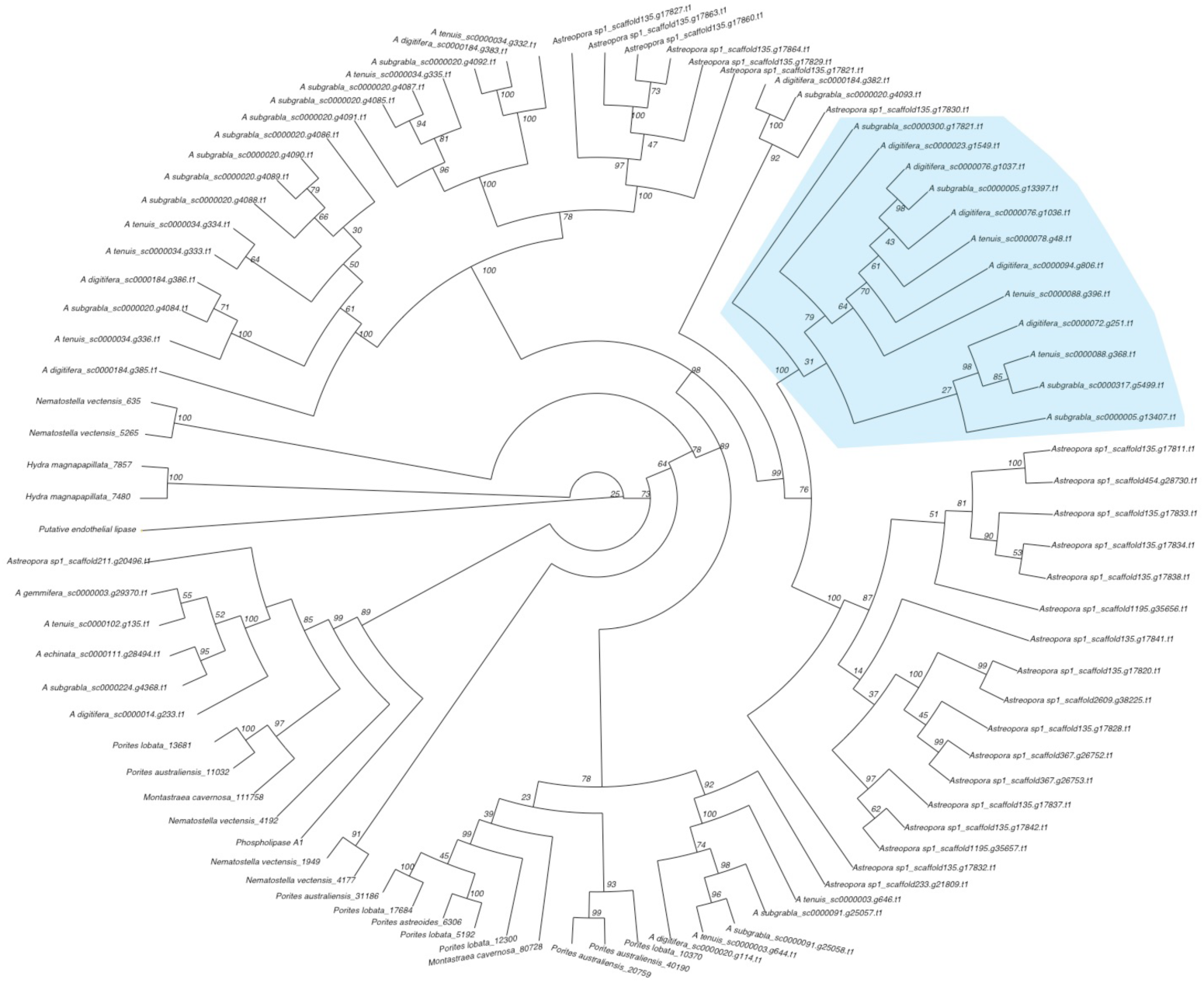
Phylogeny of the toxic putative endothelial lipase. The phylogeny was reconstructed using RAxML and bootstrap values are shown at each node. WGD in *Acropora* is shown in cyan.

**Figure S13.**
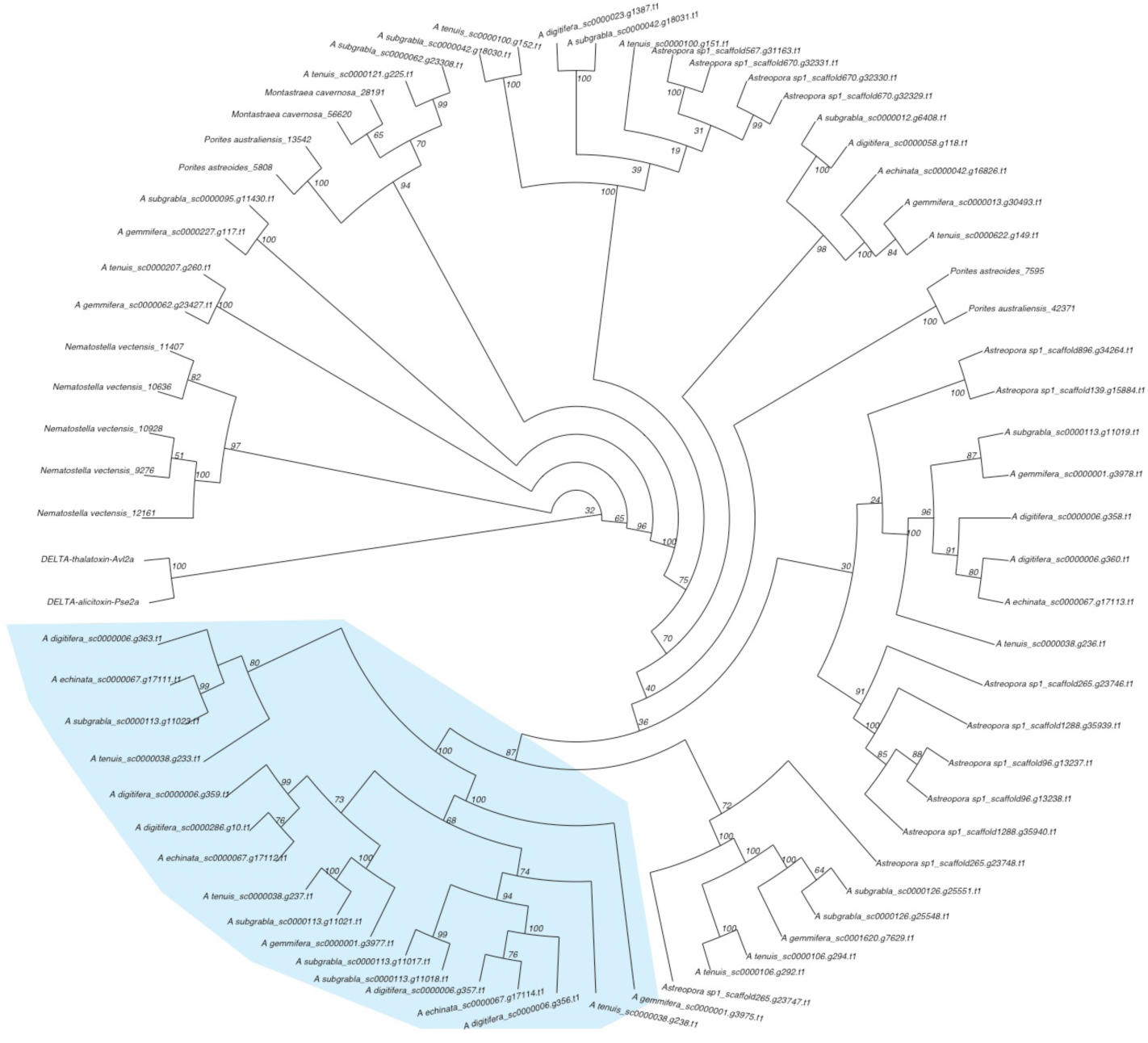
Phylogeny of the toxin protein (DELTA-thalatoxin-Avl2a/DELTA-alicitoxin-Pse2a). The phylogeny was reconstructed using RAxML and bootstrap values are shown at each node. Gene duplication by WGD in *Acropora* is shown in cyan.

**Figure S14.**
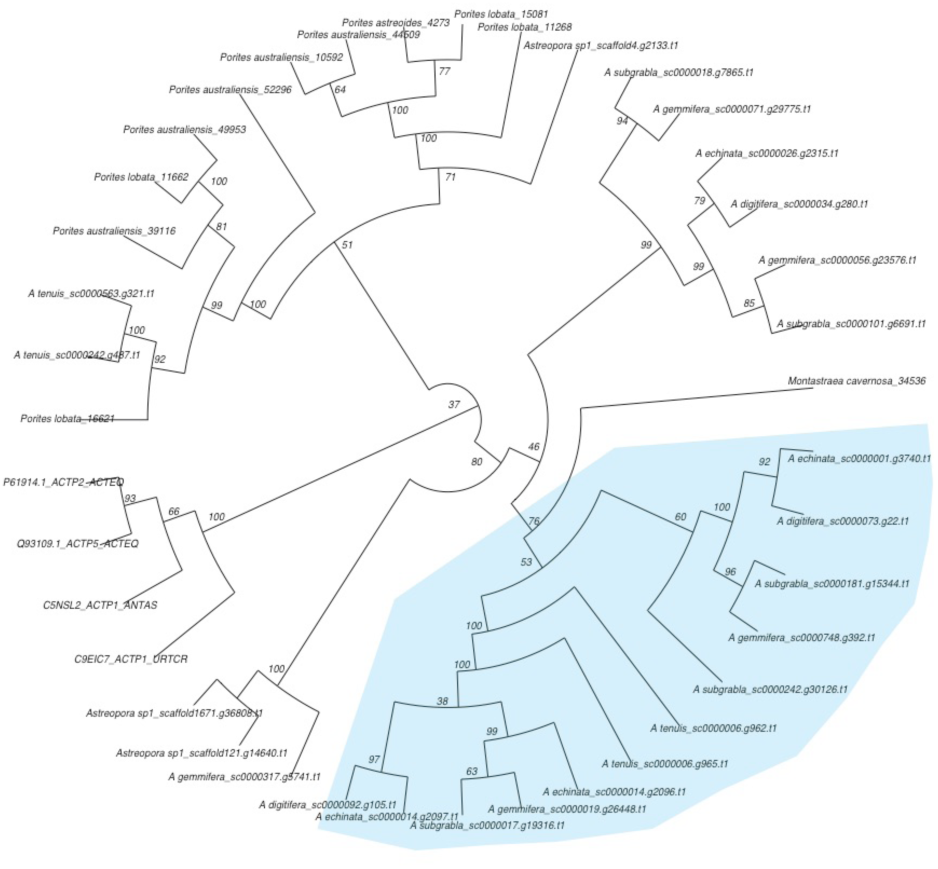
Phylogeny of the toxic protein (DELTA-actitoxin-Aas1a). The phylogeny was reconstructed using RAxML and bootstrap values are shown at each node. Gene duplication by WGD in *Acropora* is shown in cyan.

**Figure S15.**
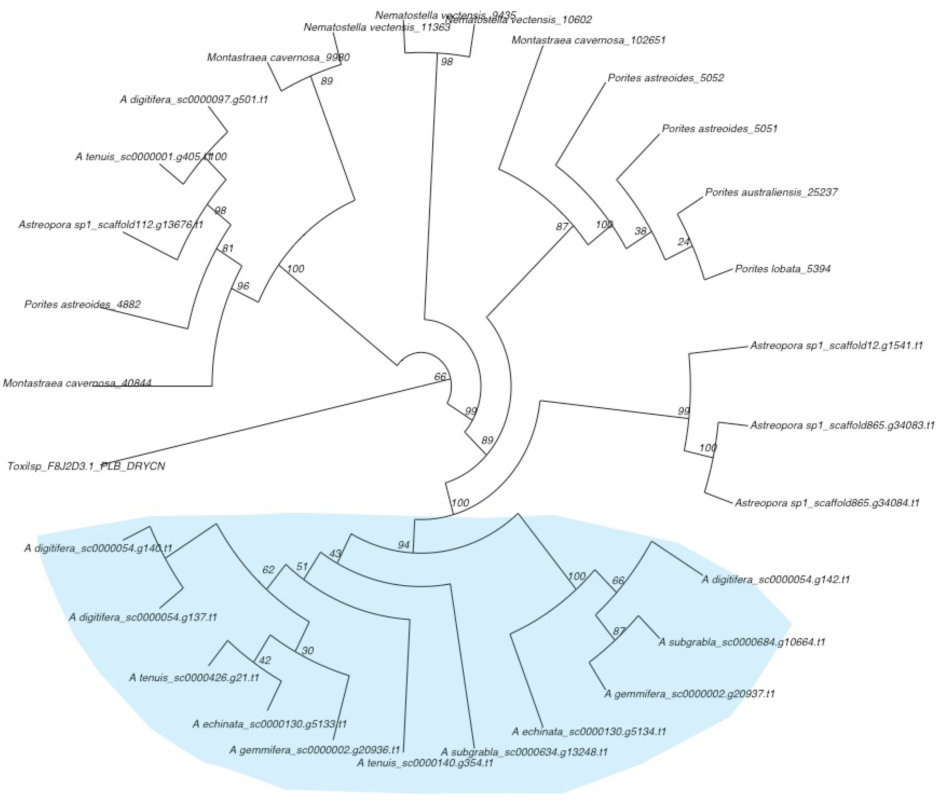
Phylogeny of the toxic protein (Phospholipase-B 81). The phylogeny was reconstructed using RAxML and bootstrap values are shown at each node. Gene duplication by WGD in *Acropora* is shown in cyan.

**Figure S16.**
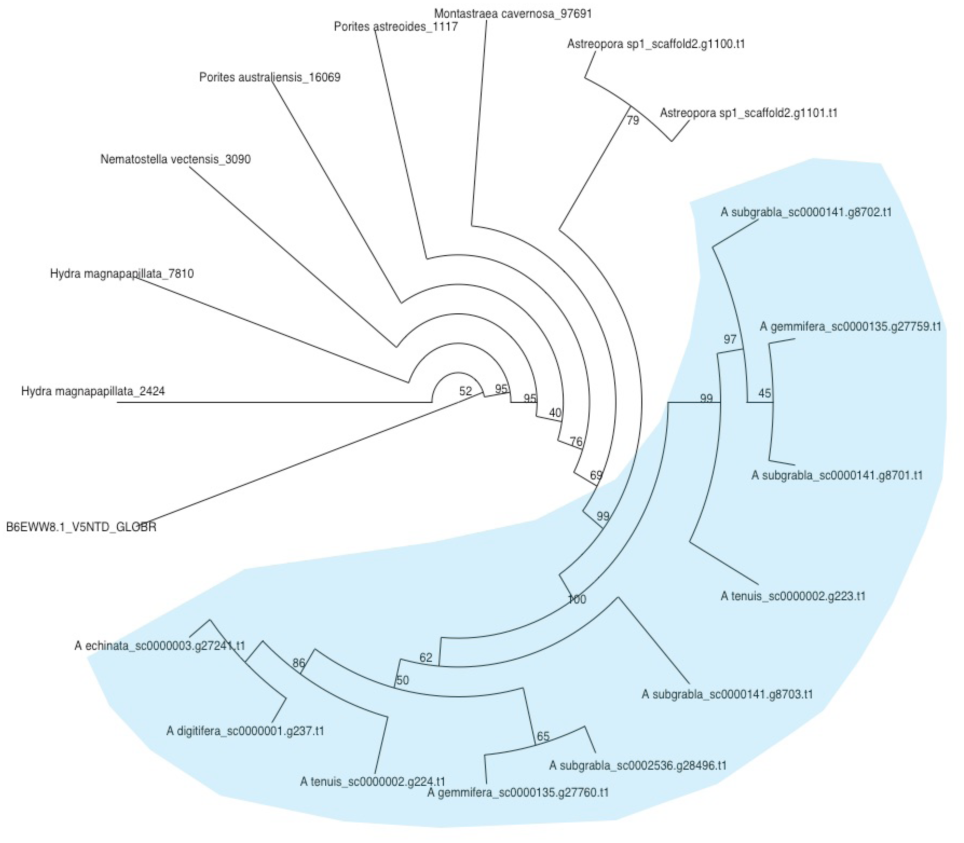
Phylogeny of the toxic snake venom 5’-nucleotidase. The phylogeny was reconstructed using RAxML and bootstrap values are shown at each node. Gene duplication by WGD in *Acropora* is shown in cyan.

**Figure S17.**
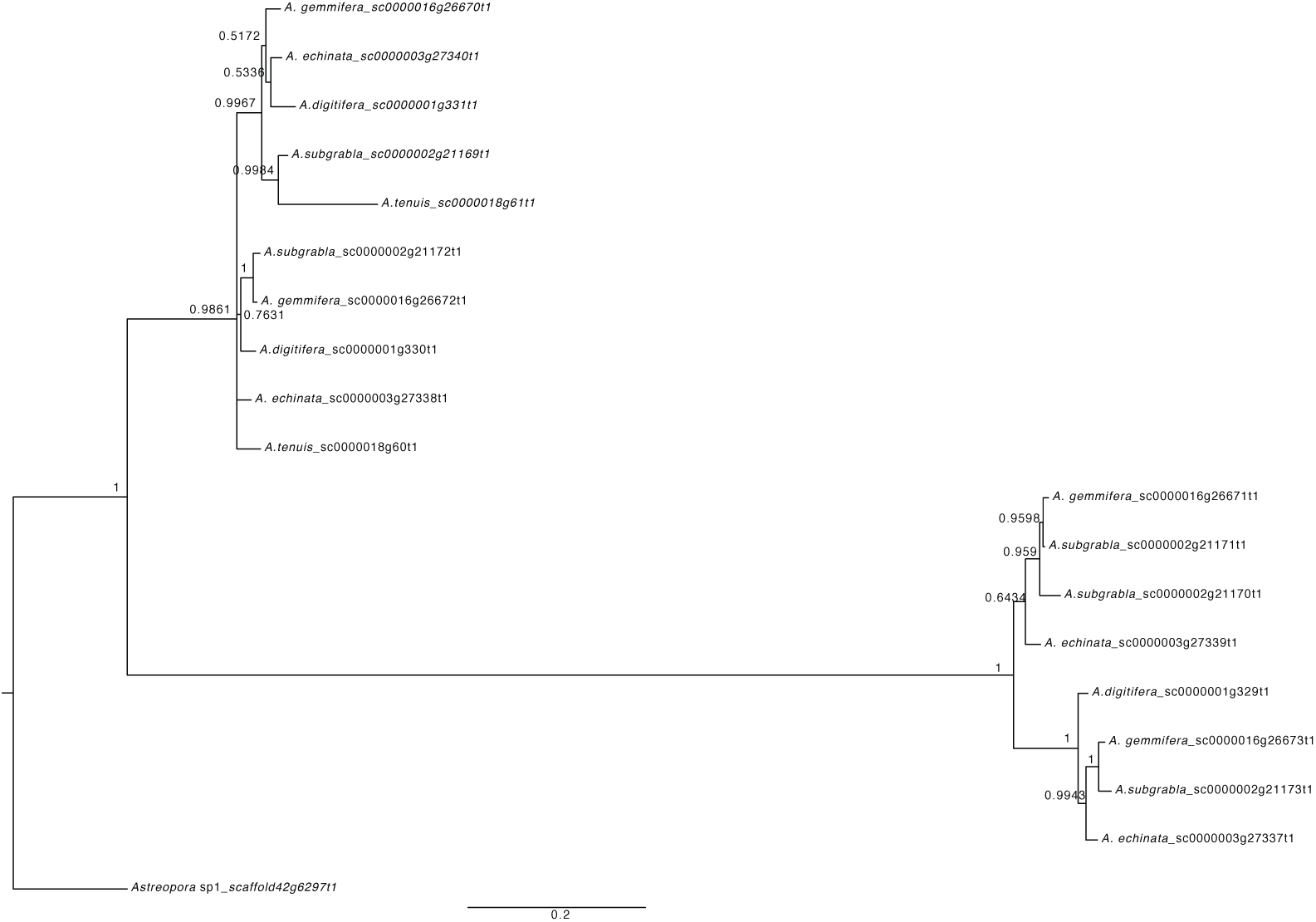
Phylogeny of orthogroup 434 (somatostatin receptor type 5-like) shows duplicates under two WGD topology. The phylogeny was reconstructed using MrBayes, and Bayesian posterior probabilities are shown at each node.

### Supplementary Tables

**Table S1.** Numbers of gene pairs in the paralogous gene pairs dataset and anchor gene pairs datasets.

| | Paragous gene pairs ($\leq 20$ gene families) | | Anchor gene pairs ($\leq 20$ gene families) | |
| --- | --- | --- | --- | --- |
| | Total numbers | Total numbers ($0 < dS < 2$) | Total numbers | Total numbers ($0 < dS < 2$) |
| *A. digitifera* | 39827 | 8249 | 46559 | 1958 |
| *A. echinata* | 47299 | 10948 | 54956 | 2530 |
| *A. gemmifera* | 48051 | 11093 | 56972 | 3299 |
| *A. subgrabla* | 50852 | 12077 | 44093 | 2380 |
| *A. tenuis* | 34097 | 6635 | 28073 | 1488 |
| *Astreopora*<br>sp1 | 49135 | 13648 | 52481 | 3033 |

**Table S2.** Peak value estimation of dS distribution by KDE toolbox.

|  | *Astreopora* sp1_*A. tenuis* | *A. tenuis*_ *A. digitifera* | *Acropora*_WGD |
| --- | --- | --- | --- |
| Peak age | 53.6 | 14.69 | 35.01458704 |
| log2(dS_paralog_peak) | -0.314 | -3.4596 | -1.8165 |
| 95%_HDP_log2(dS_paralog_peak) | (-0.22031,-0.33195) | (-3.4008,-3.5141) | (-1.7606,-2.1261) |
| 95%_HDP_Age | NA | NA | (31.18,35.7) |

**Table S3.**
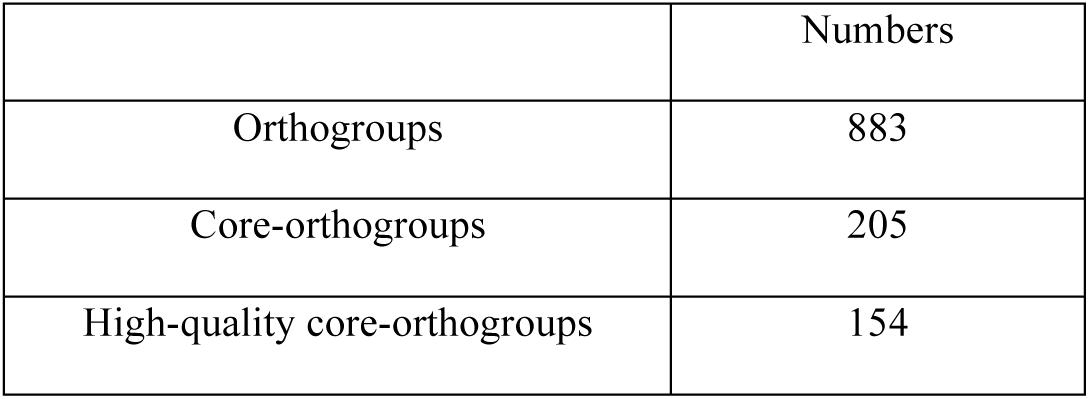
Numbers of gene family in orthogroups, core-orthogroups and high-quality core-orthogroups

|  | Numbers |
| --- | --- |
| Orthogroups | 883 |
| Core-orthogroups | 205 |
| High-quality core-orthogroups | 154 |

**Table S4.**
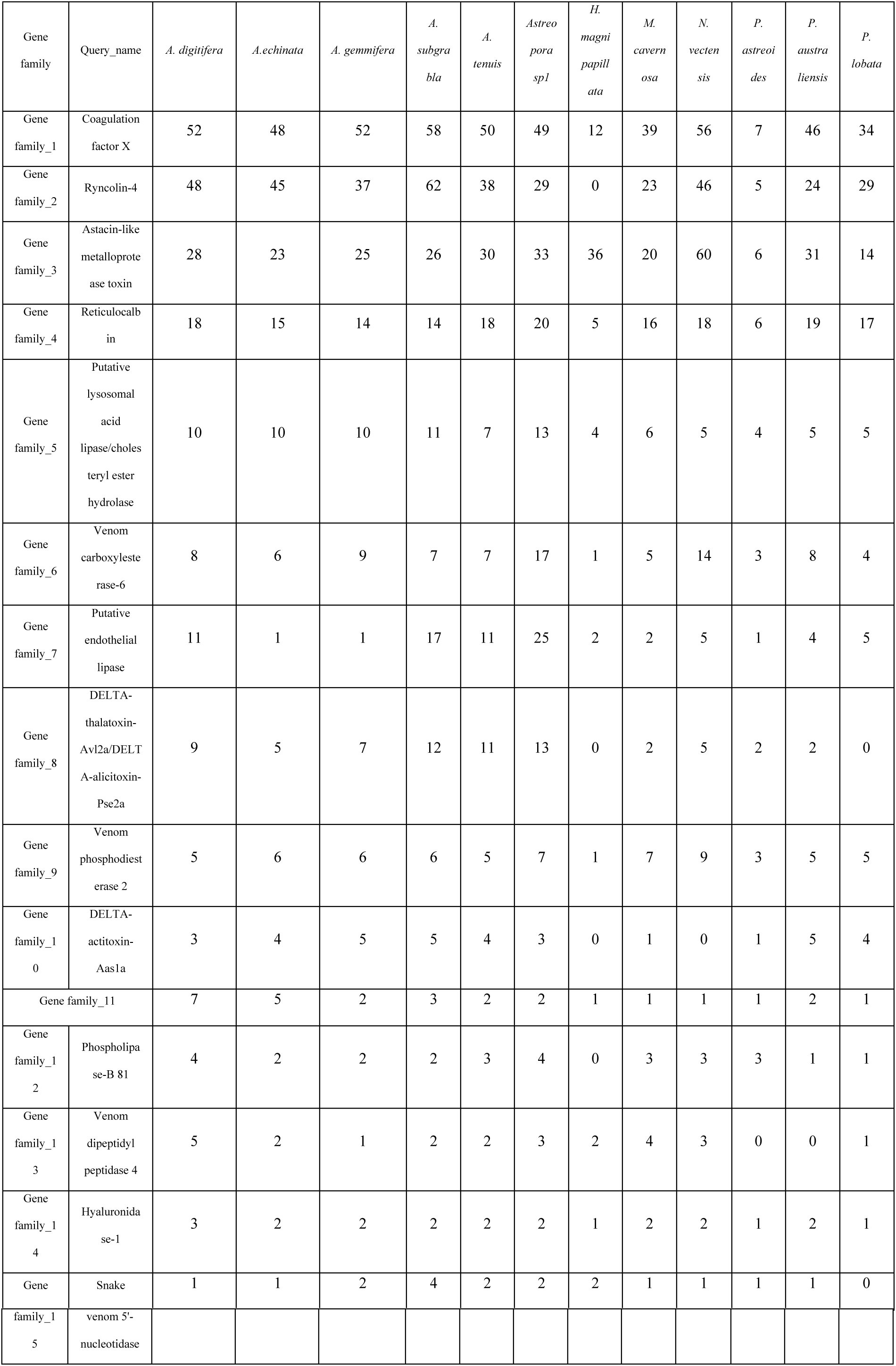
The number of putative toxin proteins in 12 Cnidarian species

**Table S5.** Likelihood of multiple WGDs hypotheses in *Acropora* using WGDgc method with gene counts data.

| WGD event(s) | Likelihood | Likelihood Ratio Test | P_value |
| --- | --- | --- | --- |
| 0 | -38731.86 | 0 VS 1 | 7.66E-05 |
| 1 | -38724.04 | 1 VS 2 | 0.01248965 |
| 2 | -38720.92 | 2 VS 3 | 0.05990546 |
| 3 | -38719.15 |  |  |

**Table S6.** Likelihood of different times of WGD under one WGD event in *Acropora* using WGDgc.

| Time of WGD | Likelihood |
| --- | --- |
| 18.697005 | -38724.14 |
| 22.697005 | -38724.09 |
| 26.697005 | -38724.06 |
| 30.697005 | -38724.04 |
| 34.697005 | -38724.04 |
| 38.697005 | -38724.06 |
| 42.697005 | -38724.11 |
| 46.697005 | -38724.2 |
| 50.697005 | -38724.35 |

